# Translocations can drive expression changes of multiple genes in regulons covering entire chromosome arms

**DOI:** 10.1101/2024.10.07.616925

**Authors:** Anna Oncins, Roser Zaurin, Houyem Toukabri, Kimberly Quililan, José R Hernández Mora, Magdalena A Karpinska, Erik Wernersson, Alastair Smith, Agostina Bianchi, Leone Albinati, Andrea Rivero, François Serra, Raúl Gómez, Cristina López, Sílvia Beà, Nadia Halidi, Alfonso Valencia, Magda Bienko, A Marieke Oudelaar, Renée Beekman

## Abstract

Translocations have largely been implicated in tumor development. However, beyond the consequences of aberrant gene expression near the breakpoint, their effects remain unexplored. In this work, we characterize the interplay between translocations, chromatin organization and gene expression using mantle cell lymphoma (MCL) as a model. We show that *in vitro* induced MCL-associated translocations can drive transcriptional changes at entire chromosome arms affecting multiple genes in a regulon-like fashion. Additionally, overexpressed genes in MCL patients are enriched in the exact same genomic regions, further underlining its potential relevance for lymphomagenesis. Moreover, we demonstrate a clear link between the translocation-induced transcriptional alterations and genome organization, with genes most susceptible to change expression residing in pre-existing long-range interacting loops spanning 50 megabases. The translocation places the strong immunoglobulin enhancer into this loop, allowing the spread of its regulatory potential over the entire affected chromosome arm. Finally, we show that translocation-induced effects mainly represent expression enhancement of genes already active prior to translocation formation, highlighting the importance of the epigenetic state of the cell in which this initial hit occurs. In summary, we show that translocations can induce ample, simultaneous gene expression changes affecting entire chromosome arms, representing an important new mechanism for tumorigenesis.

## Introduction

With the advancement of chromosome conformation capture (3C) and next-generation sequencing technologies, our knowledge of the cell’s nuclear structure has evolved drastically. It is nowadays commonly accepted that the large complexity of mammalian organisms cannot be explained only by the number of genes, nor by the sole effect of the linear genomic sequence^1,2^. In contrast, it is widely established that mammalian nuclei are highly structured organelles at the three-dimensional (3D) level, displaying different scales of increasing complexity^3^. From promoter-enhancer loops^4^ to topologically associating domains (TADs)^5^ and A and B compartments^6^, the genome is organized in a non-random fashion, contributing to gene expression variability and cell diversity. At the largest scale and beyond these intrachromosomal genome organization characteristics, a clear level of interchromosomal organization exists, whereby chromosomes occupy separate spaces, known as chromosome territories (CT) that intermingle little with each other^7^. Recent studies have furthermore shown a correlation between the radial positioning of CTs and certain chromosomal characteristics, such as length, gene density and GC content^8,9^, indicating that genome organization is neither random at the interchromosomal level.

Structural variants (SVs) including deletions, duplications, inversions, insertions and translocations, lead to a global reorganization of the genome, not only affecting its linear sequence but also 3D genome organization at all the aforementioned levels. While structural genomic rearrangements contribute to genetic diversity in eukaryotes, they can also play a key role in disease development and progression. With respect to tumor formation, since the description of the translocation t(9;22)(q34.1;q11.2), the Philadelphia chromosome, in chronic myeloid leukemia (CML) in the 1960s^10,11^, SVs have been recurrently associated with malignant transformation. For example, another translocation was discovered in acute promyelocytic leukemia (APL), in which patients often display the translocation t(15;17)(22q;q12)^12^. First attempts to understand the oncogenic effect of these SVs revealed the formation of chimeric or fusion proteins such as BCR::ABL and PML::RAR, which were generated as a direct consequence of the translocation in CML and APL respectively^12,13^. Later, new evidence showed that the effect of SVs could also rely on the formation of new intrachromosomal interactions, driving gene expression alterations beyond fusion proteins. Efforts focused then on understanding SV-induced enhancer hijacking and enhancer activation loss, for example at the *EVI1/GATA2* locus in acute myeloid leukemias (AML) carrying inversions or translocations of chromosome 3^14^. Yet, the effect of SVs, and translocations in particular, at the long-range intrachromosomal and interchromosomal level remains less well investigated. It was not until the last decade that first evidence linking SVs with global genome reorganization was provided. For instance, studies carried out on Ewing’s sarcoma patients demonstrated a significant relocalization of translocated chromosomes 11 and 22 within the cell nucleus^15,16^. More recently, the description of systematic changes of contact frequencies between short and long chromosomes in the presence of aneuploidies, such as in Patau syndrome (trisomy of chromosome 13) or Edwards syndrome (trisomy of chromosome 18), confirmed the link between SVs and nuclear reorganization^17^. Yet, the mechanisms by which SVs affect nuclear architecture and which functional consequences these changes have remain largely unclear.

In this context, non-Hodgkin lymphomas (NHLs) represent a good model to study the effect of SVs on tumor progression, since in many cases their main hallmark aberration is a translocation involving the IGH enhancer locus and different proto-oncogene harboring regions^18^. In this study, we specifically focus on mantle cell lymphoma (MCL), a lymphoid malignancy that accounts for 5-10% of all NHLs^19–21^. MCLs frequently display the t(11;14)(q13;q32) translocation, which leads to the juxtaposition of the IGH enhancer next to the *CCND1* proto-oncogene, a positive regulator of cell cycle. Nevertheless, CCND1 overexpression alone is not sufficient to drive tumor formation^22,23^, suggesting that the effect of the translocation could be broader than previously appreciated. Recent advances in chromatin organization have shed light on how enhancer-promoter interactions and A-B compartments are altered in MCL^24^. However, the effect of translocations on the global interchromosomal interaction landscape, as well as on the long-range intrachromosomal 3D organization in MCL remains largely unclear. Additionally, the consequences of these changes for lymphoma initiation and progression are still unknown. Therefore, in this study we aimed to explore the intricate interplay between translocations, chromatin architecture and gene expression, using MCL as a model. To that end, we combined chromosome conformation capture, *in silico* 3D modeling, and 3D fluorescence in situ hybridization (FISH) data from MCL patient samples and/or cell lines-allele-specific where possible - with gene expression analysis upon *in vitro* translocation generation and in MCL patient samples. Through these complementary analyses, we show that translocated chromosomes can undergo extensive genome organization alterations that are tightly linked to large-scale gene expression changes. More specifically, we describe that IGH translocations can drive gene expression changes of multiple genes covering entire chromosome arms, turning these long genomic segments into large regulatory units. Interestingly, the overexpressed genes in our *in vitro* translocation-induced models and MCL patients are enriched in the exact same genomic regions, underlining the potential relevance of our finding for lymphomagenesis. Moreover, we show that pre-existing chromatin loops are exploited by translocations, allowing to spread the regulatory effect of strong enhancers over more than 50 megabases. We furthermore demonstrate that the translocation-induced effects mainly represent an enhanced expression of genes already active prior to translocation formation. Overall, this highlights the importance of the epigenetic state of the tumor cell of origin in which this initial hit occurs. In summary, our findings can have a major impact on the early stages of tumor formation, representing an important new mechanism for tumorigenesis.

## Results

### MCL cells undergo restructuring of the interchromosomal interaction landscape

MCL cells are malignant B cells that carry a hallmark translocation involving the frequently balanced exchange of DNA between chromosomes 11 and 14, leading to the juxtaposition of the *CCND1* proto-oncogene with the IGH enhancer. While the effect of the translocation resulting in CCND1 overexpression is widely reported, other possible consequences of the translocation remain unclear. For instance, the transfer of chromosomal material leads to the formation of two new derivative chromosomes, with altered length, gene density and GC content (fig 1a). These variables have previously been associated with the spatial positioning of chromosomes within the cell nucleus^8,25–27^, suggesting that translocations possibly affect chromosome territory (CT) positioning.

**Figure 1.**
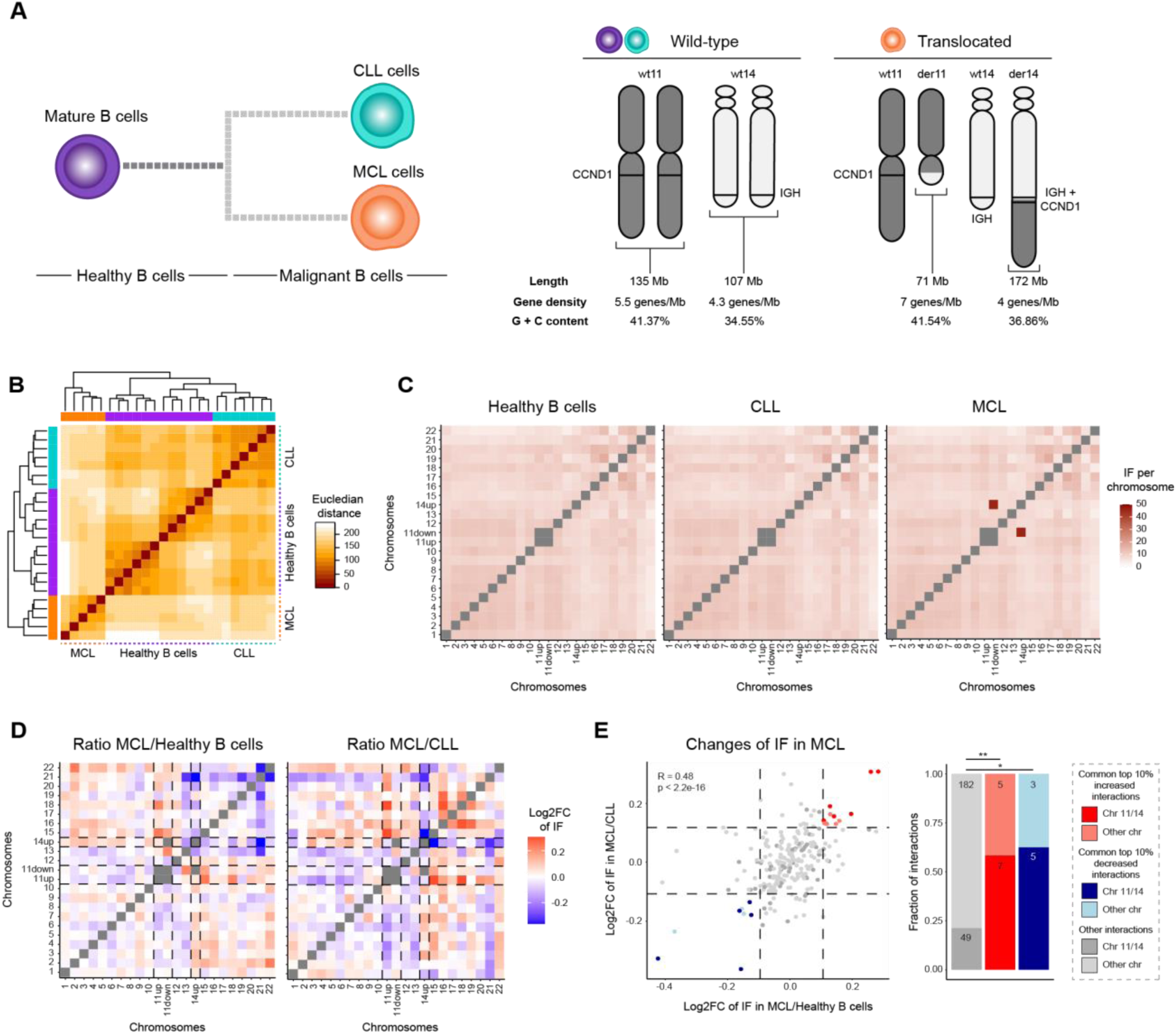
The interchromosomal 3D landscape in MCL. **A)** MCL and CLL arise from healthy mature B cells upon malignant transformation (left), with the translocation t(11;14) in MCL resulting in changes in chromosomal characteristics such as length, gene density and GC content (right). **B)** Dissimilarity matrix comparing the Hi-C matrices of all interchromosomal reads among samples. **C)** Heatmap of interchromosomal interaction frequencies between chromosomes after ICE-normalization. Chromosomes 11 and 14 are divided into “up” and “down” regions, according to their breakpoints in MCL samples. **D)** Log2 fold-change of the ratio of interaction frequencies between MCL and healthy B cells (left) and MCL and CLL (right). Dashed lines highlight translocated chromosomes, again divided in “up” and “down” as in C. **E)** Log2 fold-change of interchromosomal interaction frequencies in MCL versus healthy B cells (x-axis) or CLL (y-axis) with the top 10% common increased/decreased interaction frequencies shown in red/blue (left). Barplots of fractions of increased (red), decreased (blue) and non-changed (grey) interactions in chromosomes 11 and 14 (dark) or other chromosomes (light) (right). Numbers in the bar plots represent the total number of pairwise interactions in the given segment. der = derivative, wt = wild-type, down = downstream of the breakpoint position, up = upstream of the breakpoint position, IF = interaction frequency, **p-value < 0.01, * p-value < 0.05.

To understand the effect of the MCL-related structural variant t(11;14) on nuclear architecture, we first analyzed publicly available *in situ* Hi-C data of five MCL patients at diagnosis, and compared their interchromosomal interaction frequency (IF) landscape to mature B cells from healthy individuals as negative controls^24^ (supp table 1). The 12 control samples comprised healthy naive, germinal center and memory B cells, as well as plasma cells. Additionally, we compared the alterations in MCL to seven chronic lymphocytic leukemia (CLL) samples - mature B-cell tumors without translocation - to differentiate the effects of malignant transformation from those associated with the translocation *per se* (fig 1a). Analysis of the interchromosomal interactions at the megabase (Mb) bin scale - either including (fig 1b) or excluding (supp fig 1a) the interactions between translocation partners - revealed that MCL samples cluster separately from healthy B cells and CLL. Next, we grouped the interchromosomal interactions per sample type, thus masking donor-specific effects (supp table 2) and leaving only common effects in play. Indeed, chromosome-wise comparisons showed a high, MCL-specific interaction frequency between the lower part of chromosome 11 (chr11 down, downstream of the translocation breakpoint) and the upper part of chromosome 14 (chr14 up, upstream of the translocation breakpoint), confirming the presence of the t(11;14) translocation as the only common structural variant (fig 1c). Of note, the two Mb bins downstream of the translocation breakpoint on chromosome 14 (chr14 down) - located on der11 upon the translocation - showed low numbers of 3D interactions likely due to poor mappability, and were therefore removed from the analysis. As a consequence, interaction patterns representing der11 (between chr11 up, upstream of the translocation breakpoint and chr14 down) could not be observed in our analysis.

Next, after removing the intrachromosomal interactions between translocation partners, we calculated the pairwise chromosomal interaction changes among MCL, healthy B cells and CLL (fig 1d, supp fig 1b). Our results unveiled consistent MCL-specific interchromosomal interaction alterations involving chromosomes 11 and 14. Particularly, within the common top 10% of alterations in MCL compared to healthy B cells and CLL, interactions involving chromosomes11 and 14 were significantly enriched with respect to the remaining interactions (2.75-fold enrichment for commonly increased interactions and 2.95-fold enrichment for commonly decreased interactions, with respective chi-squared p-values 0.0086 and 0.0206) (fig 1e). Interactions from chr11 up particularly increased towards short chromosomes, mainly chromosomes 15 and 18, while they decreased to long chromosomes, such as chromosomes 6 and 7. In contrast, increased interactions from chr14 up mainly occurred towards long chromosomes, such as chromosomes 3 and 7, while decreasing interactions to short chromosomes like chromosomes 15, 20 and 21 (supp table 3). From these results, we conclude that MCL cells display global alterations of the interchromosomal 3D interaction landscape, mainly associated with the chromosomes that are directly affected by the translocation.

### Derivative 11 shifts towards the nuclear center in MCL cells

In diploid cells, Hi-C analyses elucidate mean interaction frequencies of the two homologous chromosomes. Hence, allele-specific analyses are crucial to discern whether the MCL-specific interchromosomal interaction differences that we detect for chromosomes 11 and 14 originate from the translocated allele or its wild-type counterpart. To obtain these insights, we performed Hi-C in the MCL line Z-138, which carries the t(11;14) translocation, and simultaneously established bivariate flow cell karyotyping to isolate DNA from its derivative 11, wild-type 11 and wild-type 14 chromosomes (supp fig 2a). The difference in length and GC content of the derivative 11 compared to the wild-type chromosome (fig 1a) leads to the formation of an aberrant chromosome cloud in the bivariate flow cell karyotyping, which was not detected in the control cell line, and could therefore be isolated. Unfortunately, we could not detect the cloud belonging to derivative 14, possibly due to the fact that its length and GC content are similar to other chromosomes, avoiding a clear separation. Consequent sequencing of the isolated chromosomes in combination with heterozygous SNP assignment, aided by previously published array data^28^, allowed us to genotype the derivative and wild-type alleles in an allele-specific manner, enabling the classification of Hi-C reads belonging to the wild-type or the derivative alleles in Z-138 (fig 2a, supp fig 2b, supp table 4 and 5).

**Figure 2.**
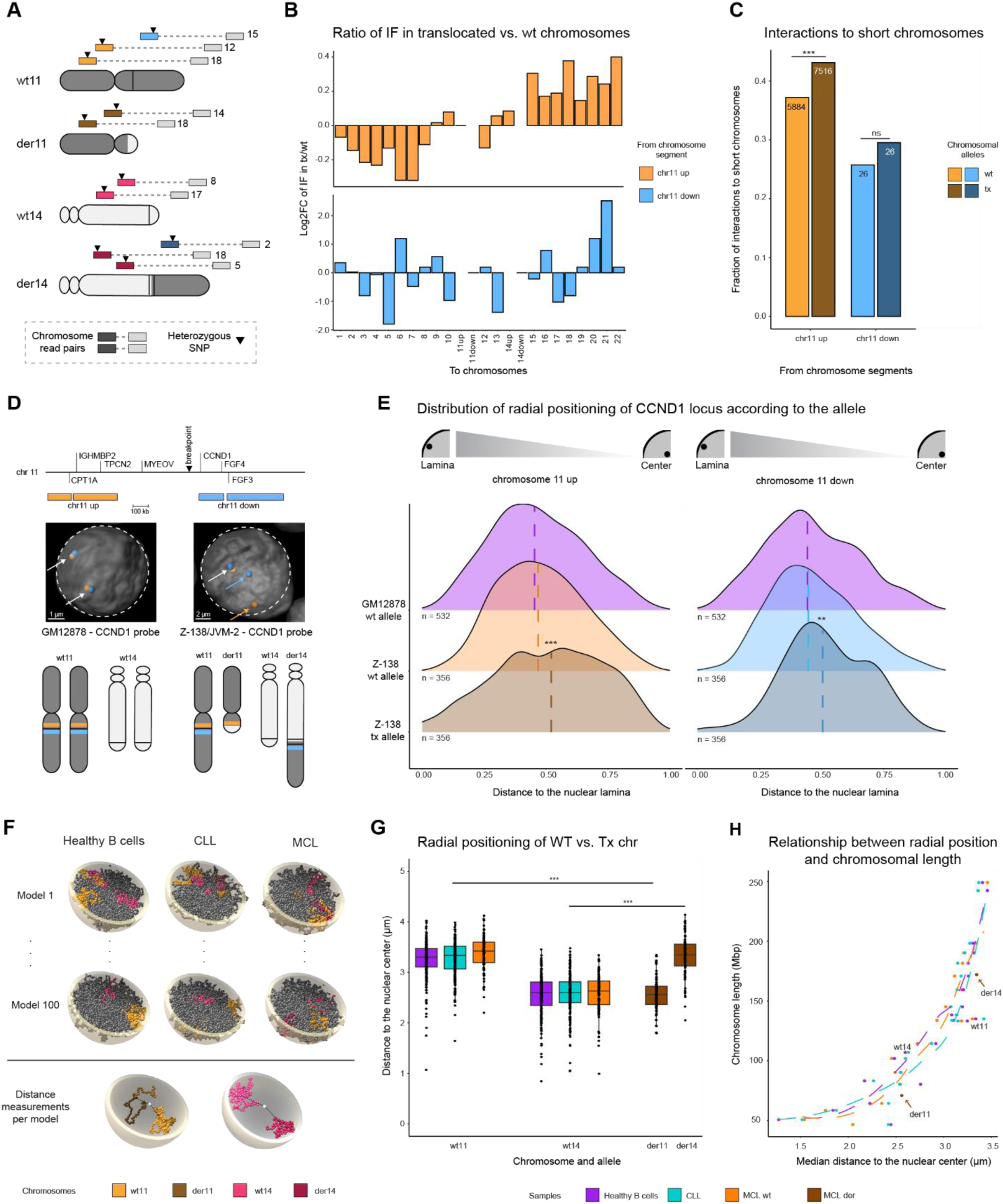
Allele-specific analysis of translocated chromosomes in MCL. **A)** Classification of Hi-C reads to the wild-type or translocated allele in Z-138. Different colored bars represent Hi-C reads from wt11 up or tx11 up (light and dark orange), wt11 down or tx11 down (light and dark blue) or wt14 up or tx14 up (light and dark pink) to other chromosomes (grey bars, hypothetical chromosome numbers are indicated) linked via the dashed line. **B)** Log2FC of the ratio of Hi-C interactions between translocated and wild-type chr11 up (upper panel) and chr11 down (lower panel) towards all other chromosomes in Z-138. **C)** Fraction of Hi-C reads from chromosome 11 (wild-type or translocated) to short chromosomes (chromosomes 13-22) in Z-138. Numbers in the bar plots represent the total number of interactions in the given segment. **D)** Position of the *CCND1* breakapart probe in wild-type and translocated chromosome 11. Two example images were cropped from the original files. White dashed lines delimit the nuclear lamina. White arrows point to wild-type chromosomes, and blue or orange arrows indicate translocated segments of chromosome 11. **E)** Density plots of the relative distribution of imaging-based radial position of chromosome 11 in Z-138 compared to control (GM12878), 0.0 indicates the closest pixel to the lamina and 1.0 the most central position. Dashed lines represent the median. **F)** Graphical representation of two random models (out of 100), colored parts represent chromosomes 11 and 14 (wild-type and translocated). Dashed lines outline the measurements from the central mass of each chromosome to the nuclear center (lower panel). **G)** Boxplots with the *in silico* modeling-based position of wild-type or translocated chromosomes 11 and 14. Healthy B cells and CLL contain 200 measurements for each chromosome, while in MCL the two alleles are divided into 100 wild-type and 100 derivative alleles. **H)** Relationship between the *in silico* modeling-based radial position and the chromosome length of each chromosome. The length of derivatives 11 and 14 was calculated according to the breakpoint bin in MCL patients. Colored lines indicate the estimated loess curve for each sample, the curve in MCL was calculated without considering the derivatives. Brown arrows indicate the derivatives. der = derivative, wt = wild-type, tx = translocated, down = downstream of the breakpoint position, up = upstream of the breakpoint position, IF = interaction frequency, **p-value < 0.01, ***p-value < 0.001, ns = not significant.

The clear gain of interactions between the translocated chromosomes 11 and 14 in comparison to the wild-type chromosomes confirmed the correct classification of Hi-C reads (supp fig 2c). We furthermore detected an increased number of reads between the considered wild-type chromosome 14 and chromosome 8, which appeared to represent a t(8;14)(q24;q32) translocation that was previously reported^29^ (supp fig 2c). From this result, we concluded that Z-138 does not carry a wild-type copy of chromosome 14 to compare with derivative 14 in our allele-specific analyses. Therefore, chromosome 14 was excluded from further analysis in Z-138. After removing the translocation-based intrachromosomal read counts, we observed a significant increase of interactions from the translocated upper part of chromosome 11 to short chromosomes (13 to 22) in comparison to wild-type (chi-square test, p-value < 2.2e-16) (fig 2b-c), indicating a possible shift of derivative 11 towards the nuclear center. We could not detect any significant interaction changes from the translocated lower part of chromosome 11 towards long or short chromosomes. However, the low number of reads that could be classified in this analysis does not allow us to conclude that such changes do not exist (fig 2c).

Chromosome conformation capture techniques cannot assess the physical position of (wild-type and translocated) chromosomes 11 and 14 within the nucleus, as they only define the relative interaction frequency between genomic fragments. For this reason, we performed allele-specific high-throughput FISH in Z-138 cells, using break-apart probes that specifically bind upstream or downstream of the translocation breakpoint on chromosome 11 (fig 2d). This approach enabled us to differentiate the wild-type chromosome 11 from its translocated counterpart, and to compare their radial position within the nucleus (fig 2e). The lymphoblastoid cell line GM12878 was used as control, while JVM-2 cells - also carrying the t(11;14) translocation - were included to account for possible variability between MCL cell lines. For each cell line, 3D images were generated, followed by assigning hybridization signals to individual cells. Next, probe pairs located up- and downstream of the breakpoint were categorized as wild-type if located close in 3D space (fig 2d), falling within the 95% confidence interval established using a model fitted in GM12878 control cells. The remaining, more distant pairs were classified as translocated. Only those cells containing one wild-type and one translocated pair (Z-138 and JVM-2) or two wild-type pairs (GM12878) were considered for further analysis. Next, the 3D distance from each probe to the nuclear surface was measured. In line with our previous Hi-C results, FISH analyses showed a significant shift of the radial position of the translocated upper part of chromosome 11 in Z-138, which was located closer to the nuclear center compared to wild-type 11 (fig 2e, left, Wilcoxon test p-value = 2.292e-05). A similar trend was observed in JVM-2 cells (supp fig 2d). Similarly, the translocated lower part of chromosome 11 displayed a significant shift towards the nuclear center in Z-138 compared to wild-type 11 (fig 2e, right, Wilcoxon test p-value = 0.01267), although this movement was not detected in JVM-2 (supp fig 2d). Additionally, our data point towards a clear bimodal trend of radial position of the translocated chr11 down segment in both Z-138 and JVM-2 cells, suggesting the possible presence of distinct subpopulations regarding its nuclear positioning. We also analyzed chr14 positioning using JVM-2 cells, but unfortunately the data was too sparse to obtain significant results, due to a high level of tetraploidy in this cell line (supp fig 2d).

To further explore the chromosomal radial distribution in the presence or absence of the t(11;14) translocation, we leveraged the patient-derived Hi-C data to generate 100 *in silico* models per sample type - MCL, CLL and healthy B cells - using Chrom3D^30^. To generate diploid cellular models we deconvoluted the Hi-C data considering that, based on our previous results, wildtype chromosomes 11 and 14 in MCL retain the interaction frequencies of normal B cells, while the translocated alleles accounted for the differences observed between MCL and normal B cells. Each model represented a possible diploid nuclear chromatin conformation structure, for which CT positioning was inferred by Chrom3D using the interaction frequencies among chromosomes in the Hi-C data (fig 2f). Next, we measured the distance from the central mass of each chromosome to the geometric nuclear center, and compared their radial distribution among MCL, CLL and healthy mature B cells. While we did not observe major differences between the wild-type chromosomes in MCL versus CLL and healthy B cells (supp fig 2e), we detected a significant shift in the position of the translocation partners (Student’s T-test, p-values < 2.22e-16) compared to wild-type chromosomes 11 and 14 (fig 2g). More specifically, we noticed a median inward movement of derivative 11 in MCL over 17.2% of the radial distance (0.86/5 µm, fig 2g), displaying a similar nuclear location as wild-type chromosome 14. On the other hand, derivative 14 shifted outwards over a distance covering 14.2% of the radial length (0.71/5 µm), taking a radial position similar to wild-type chromosome 11. These results pinpoint a possible exchange of the radial position of the translocation partners within the nucleus, which aligns with the expected position based on their new length (fig 2h).

Altogether, the combination of our allele-specific Hi-C, imaging and modeling analyses indicate a shift of derivative 11 towards the nuclear center in comparison to wild-type chromosome 11, strengthening our observations of the MCL patient Hi-C analysis (fig 1c). Furthermore, our modeling data suggests a shift of derivative 14 towards the nuclear periphery compared to wild-type chromosome 14, although we do not have sufficient imaging data, nor allele-specific Hi-C data to prove this finding. Therefore, we refrain from drawing this conclusion as we consider that further studies are needed to confirm the nuclear positioning of derivative 14.

### Derivative 14 displays new long-range interactions with increased regulatory potential

Our previous observations indicate that the derivative chromosomes formed upon the t(11;14) translocation in MCL cells exhibit the most prominent interchromosomal structural alterations compared to wild-type cells. Therefore, we next investigated whether these effects extend to intrachromosomal interactions, particularly focusing on differences between the derivatives and their wild-type counterparts, and on the establishment of new long-range interactions. To that end, we first leveraged our MCL, CLL and healthy B-cell *in silico* nuclear models (fig 2f) to calculate the distances among all Mb bins on chromosomes 11 and 14 (fig 3a). The comparison between the wild-type and translocated chromosomes revealed a global shift in intrachromosomal distances, predominantly affecting the telomeres and the breakpoint regions on chr11 and chr14 up in an opposite manner (fig 3b). Of note, while the breakpoint region on chr11 up gains linear proximity to the telomere upon translocation t(11;14), this proximity is lost at the breakpoint region of chr14 up (fig 1a). Our data suggests that the gain or loss of linear proximity to the telomere results in a respectively larger (chr11 up, fig3b, left) or smaller (chr14 up, fig3b, right) distance between the bins located close to the breakpoint and the middle segment of the chromosome in 3D space. We thus hypothesize that telomeric displacement affects global chromosome folding, whereby 3D proximity between telomeres and genomic segments in the middle of the chromosome are disfavoured. We furthermore observed focal gains of proximity downstream of the breakpoint within derivative 14, suggesting stronger loop formation and/or new long-range interactions within this segment upon translocation. To get a more detailed view of this region, we investigated the 3D distance of all bins on chr11 down to the breakpoint. From this analysis, we observed that healthy B cells and MCL cells share a region on chr11 down, around Mb bin 120, that is especially close to the translocation breakpoint in 3D space. Interestingly, based on this pre-existing, 50 Mb spanning long-range loop in healthy B cells, the region around Mb bin 120 on chr11 down gets close to the strong IGH enhancer in 3D space upon t(11;14) translocation formation (fig 3c). Thus, while translocations do not necessarily alter long-range chromosomal loop structures they can instead exploit pre-existing loops to establish new 3D interactions with potentially strong regulatory impact.

**Figure 3.**
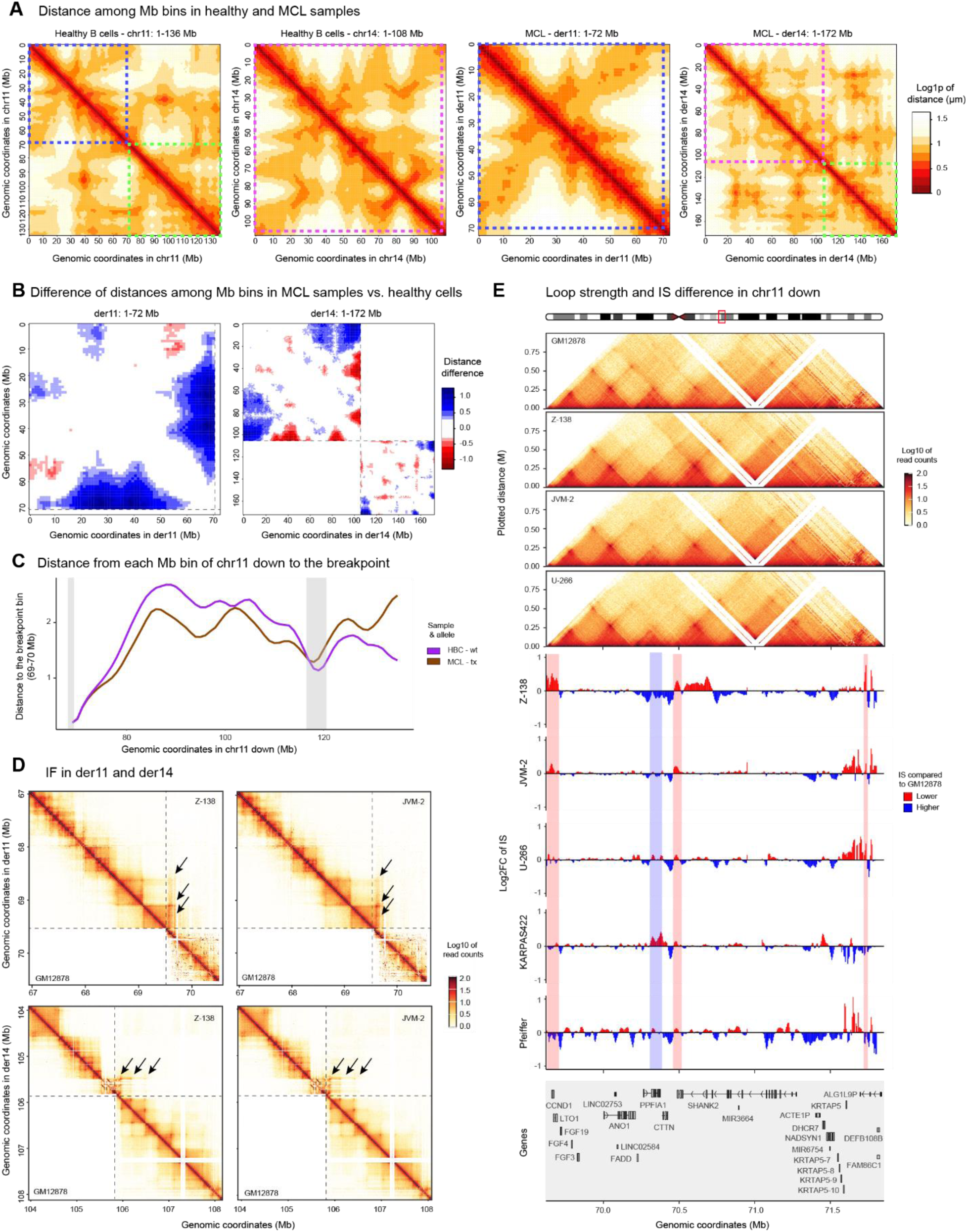
Intrachromosomal interaction landscape of translocated chromosomes in MCL. **A)** Log1p-transformed distance in micrometers between the position of each megabase bin in chromosomes 11 and 14 (wild-type, in healthy B cells, left 2 panels) and derivatives 11 and 14 (in MCL, right two panels). Dashed boxes indicate the chromosomal segments up- and downstream of the translocation breakpoint in t(11;14). Distances were inferred from the chrom3D modeling output. **B)** Differences between the position of each Mb bin in MCL compared to healthy B cells. Blue indicates a higher difference, these bins are thus located further away in MCL than in healthy B cells. On the contrary, red indicates a smaller distance in MCL compared to healthy B cells. Dashed lines indicate the translocation breakpoint. **C)** Distance from each Mb bin to the translocation breakpoint at the *CCND1* locus (Mb bin 69-70). Grey bars indicate regions closer to the breakpoint in 3D space in both healthy and MCL samples. **D)** Tiled-C read count heatmaps in derivatives 11 and 14 comparing Z-138 and JVM-2 cell lines carrying the t(11;14) translocation (right-upper triangle) with the control GM12878 (left-bottom triangle). Black arrows highlight interactions spanning over the breakpoint, indicated by the dashed line. **E)** Differences of insulation score on chr11 down in all analyzed cell lines compared to GM12878 cells. In KARPAS422 and Pfeiffer, chr11 down is not affected and should be considered as wildtype control. Colors indicate a higher (blue) or lower (red) insulation score. The heatmaps represent Tiled-C read counts in the represented segments. der = derivative, wt = wild-type, tx = translocated, down = downstream of the breakpoint position, up = upstream of the breakpoint position, IS = insulation score.

To better understand chromatin conformation changes taking place in regions proximal to the breakpoint, we next generated and analyzed high-resolution Tiled-C data^31,32^, in which we captured large regions (∼3Mb) surrounding the *IGH* and *CCND1* loci in different cell lines carrying either the common t(11;14) translocation (Z-138 and JVM-2) or a cryptic insertion of the IGH enhancer inside chromosome 11 (U-266), as well as in a wild-type control cell line (GM12878). In order to better characterize the effect of the IGH translocation independently of the translocation partner, we also included the analysis of two cell lines (KARPAS422 and Pfeiffer) carrying the t(14;18)(q32;q21) translocation, which is commonly found in follicular lymphoma (FL). Principal component analysis (PCA) analysis of the triplicate samples indicated that cell lines clustered based on their intrachromosomal interaction landscape (supp fig 3a). Interestingly, while the 3D genome architecture within the investigated regions on chromosomes 11, 14 and 18 was largely conserved upon translocation formation, *de novo* interactions between the translocation partners could be detected (fig 3d, supp 3b,c,d). These new interactions reached beyond the neo-TAD formed close to the breakpoint, indicating the formation of long-range interactions spanning a larger region than anticipated. Unfortunately, our captured region was too small to visualize the previously observed long-range interaction spanning 50 Mb in chr11 down. Since strong enhancers are associated with high insulated areas, and we detected stronger focal contacts in the chromosome segment that is placed in proximity of the strong IGH enhancer (chr11 down, fig 3b), we next refined our analysis by comparing insulation scores between cell lines with and without translocations. These analyses showed that derivative 14 - consisting of chr14 up, fused to chr11 down (in Z-138 and JVM-2) or chr18 down (in KARPAS422 and Pfeiffer) - displayed many subtle intrachromosomal alterations in insulation scores (fig 3e, supp fig 3e,f,g). Both chr11 down and chr18 down for example, which are juxtaposed to the IGH enhancer, alter their loop strength and insulation levels directly downstream of the breakpoint. This effect is stronger upon IGH-translocation formation than upon IGH enhancer insertion (U-266). In addition, just upstream of the breakpoint on chr14 up, we observe clear gains of insulation throughout the segment, except for the part more distal from the breakpoint, which displays stronger interactions within existing loops (supp fig 3e,g). The increased insulation score levels are also observed in U-266, in which the IGH enhancer is deleted from one of its copies of chromosome 14 (data not shown). This makes us hypothesize that the observed changes may be due to an IGH enhancer effect, but further research is necessary to better understand this phenomenon. Finally, in agreement with major effects being driven by the IGH enhancer, we do not detect any specific alteration in derivatives 11 and 18, which do not gain the IGH enhancer (data not shown).

Altogether, we show that derivative 14 undergoes a higher level of intrachromosomal chromatin architecture restructuring than derivative 11. Interestingly, this implies that the shift of derivative 11 towards the nuclear center does not drastically change intrachromosomal genome organization. An additional striking finding on chr11 down is the presence of a conserved long-range loop spanning the entire chromosome arm in healthy B cells and MCL. The translocation places the strong immunoglobulin enhancer into this loop, potentially allowing the spread of its regulatory potential over the entire affected chromosome arm, beyond the previously reported CCND1 overexpression. Therefore, shedding light on gene expression dysregulation in the genomic regions displaying conserved or newly formed loops could identify new alterations involved in MCL pathogenesis.

### Translocations induce large-scale gene expression effects on derivative 14

In order to assess the functional effects associated with the presence of the t(11;14) translocation, we next analyzed published RNA-seq data to evaluate gene expression changes in MCL patients compared to healthy B cells^24^. Surprisingly, we detected that the lower part of chromosome 11 contained the highest level of enrichment of MCL-specific upregulated protein-coding genes (FC 2, FDR < 0.1, fig 4a-c, supp table 6). Of note, in this analysis we decided to focus only on those genes expressed in more than half of the MCL patients - representing the vast majority of the upregulated genes - to elucidate global transcription alterations in the presence of the translocation, rather than potential patient-specific effects. Nevertheless, we recognize that the latter could also relate to MCL development. Next, we observed that the upregulated genes were not randomly distributed over chr11 down. As expected, a subset of upregulated genes was located close to the breakpoint (fig 4d). Interestingly however, we also detected a group of upregulated genes at a region located approximately 50Mb away from the translocation breakpoint - around Mb bin 120 - that gets in close proximity to the IGH enhancer in 3D space upon translocation formation (fig 3a,c). A permutation test confirmed that this distribution was indeed not random, showing an enrichment of MCL upregulated genes in Mb bin 118 to 120 (p-value 0.0402). This test furthermore pointed towards other bins containing a significant enrichment of upregulated genes, such as Mb bin 106-108 on chromosome 11 (p-value 0.0214). Based on these results, we hypothesize that IGH enhancer hijacking by distant genes located on chr11 down may occur due to translocation-induced long-range interactions. On the other hand, we could not identify a global enrichment of upregulated genes in chr11 and chr14 up(fig 4a,c, supp fig 4a). However, upregulated genes on chr11 up were grouped close to the telomere and at the breakpoint region (fig 4e), with Mb bin 67-69 clearly enriched for upregulated genes(p-value 0.0278). Overall, we hypothesize based on this data that the shift of derivative 11 towards the nuclear center may favor the upregulation of these genes.

**Figure 4.**
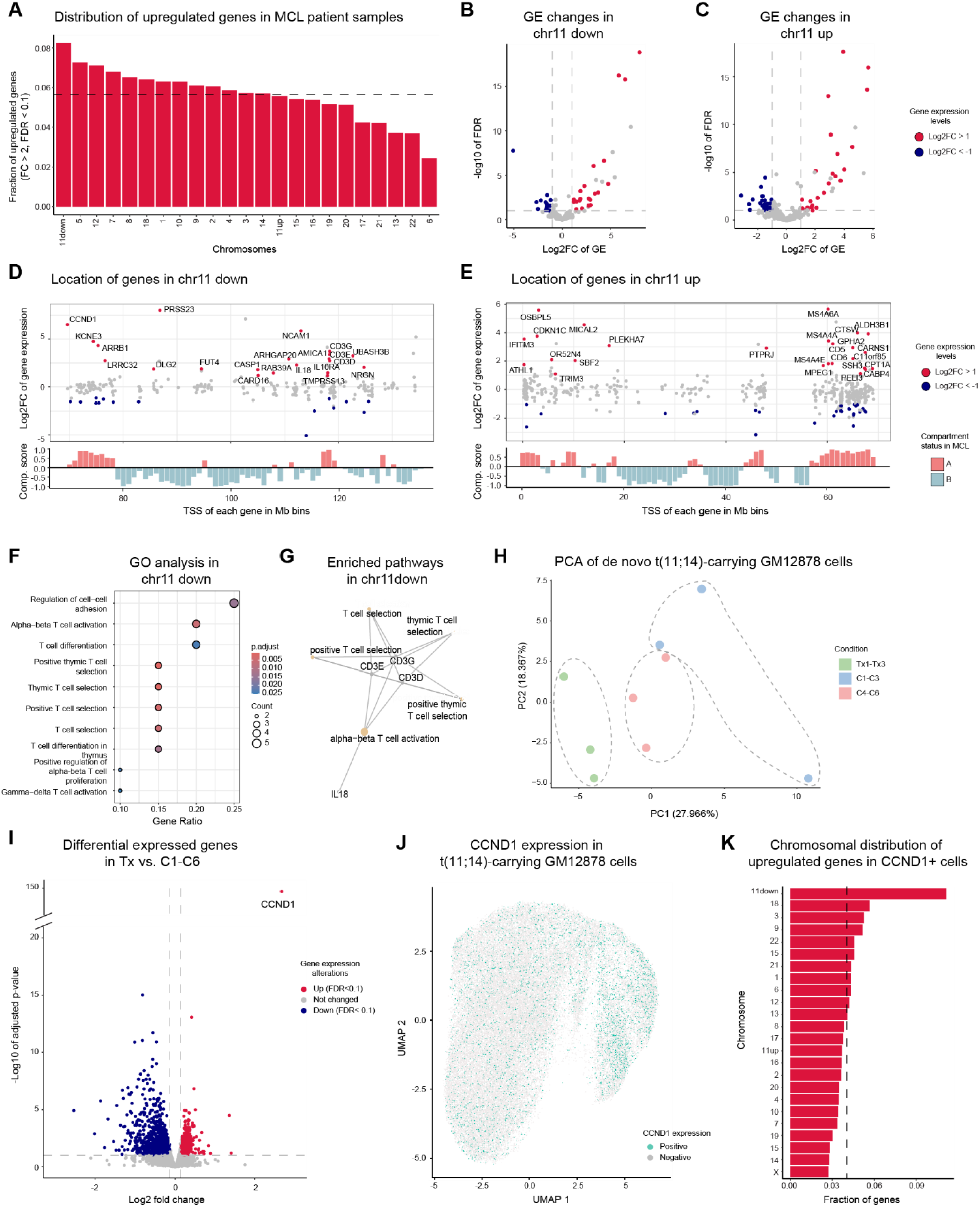
Gene expression analysis in cells carrying the t(11;14) translocation. **A)** Fraction of upregulated protein-coding genes in MCL patients versus mature healthy B cells divided per chromosome. Dashed line represents the expected fraction of genes. **B)** Volcano plot of up- and down-regulated protein-coding genes in chr11 down. Dashed lines show the absolute log2 fold change threshold of 1, and the –log10 value of the FDR value of 0.1. **C)** Same as in B, but showing up- and down-regulated genes in chr11 up. **D)** Graphic representation of the location of up- and down-regulated genes in MCL patients throughout chr11 down. Names for upregulated genes with a log2 fold change higher than 1 are depicted. Below, the compartment score of each Mb bin is represented, whereby blue represents B compartments, and red A compartments. **E)** Same as D but for chr11 up. **F)** Proportion of genes from the dataset that are associated with upregulated pathways by GO analysis. **G)** Network plot where nodes represent the genes linked to the enriched pathways depicted in F, and edges indicate the genes that are connected through the same pathways. **H)** PCA of samples carrying *in vitro* translocations or controls based on bulk RNA-seq data. **I)** Volcano plot of gene expression changes in samples carrying *in vitro* translocations or controls. **J)** UMAP of samples carrying *in vitro* translocations in a subset of their cells, based on scRNA-seq data. **K)** Fraction of upregulated protein-coding genes in CCND1-positive versus -negative cells per chromosome, after *in vitro* induction of translocations. Dashed line represents the expected fraction of genes. wt = wild-type, down = downstream of the breakpoint position, up = upstream of the breakpoint position.

Gene ontology analyses suggested a significant enrichment of T-cell-related pathways (fig 4f,g, supp fig 4b,c), as well as apoptotic- and necroptotic-related gene networks (supp fig 4b,c) in upregulated genes on chr11 down and chr14 up, respectively. Of note, Mb bin 118-120 on chromosome 11 harbors the *CD3* Epsilon, Delta and Gamma subunits, *AMICA1, IL10RA* and *TMPRSS13*. While the *CD3* genes encode for the different subunits of the T-cell receptor CD3 complex, IL10RA and TMPRSS13 are related to MAPK and TGF-beta signaling pathways, and AMICA1 controls the activation and migration of leukocytes. Gene ontology analyses of upregulated genes in chr11 up did not detect any enrichment of specific pathways. Nevertheless, we want to highlight the upregulation of CD5, MS4A paralogs and CTSW, among others, which cluster between Mb bins 60-69, close to the breakpoint region (fig 4e). CD5 is a known surface marker present on MCL cells, MS4A4 and MS4A6 are associated with macrophage-related immune responses, while CTSW regulates T-cell cytolytic activity.

The transcriptional changes described above are intriguing. However, whether they are a direct consequence of the translocation cannot be concluded from RNA-seq analysis in patient samples. In order to prove the relationship between the observed gene expression changes and the presence of the t(11;14) translocation, we generated the t(11;14) translocation *de novo* in the healthy lymphoblastoid B cell line GM12878. To that end, we nucleofected GM12878 cells with ribonucleoprotein complexes (RNPs) targeting regions on chromosomes 11 and 14 close to the translocation breakpoints described in MCL patients. As a consequence of the induced double-strand breaks, the t(11;14) translocation was formed in a subset of cells, estimated to represent 10-12% of the entire population by digital PCR (supp fig 4d). We furthermore detected an approximately 9-fold increase in CCND1 levels by qPCR in these heterogeneous populations compared to the controls (nucleofected with only one gRNA on chromosome 11 or 14), showing that the presence of 10-12% translocated cells is enough to identify strongly upregulated transcripts (supp fig 4e). Next, we performed bulk RNA-seq of the populations harboring cells with the translocation, as well as control cell populations. We could clearly distinguish them by PCA (fig 4h) and identified 1086 upregulated and 1206 downregulated genes (FC 1.1, FDR < 0.1, fig 4i, supp fig 4f, supp table 7). Although we did not see a clear enrichment of upregulated genes on chr11 down, it still harbored more upregulated genes than expected by a random distribution (supp fig 4g). Moreover, the permutation test pointed again towards a non-random distribution of bins with an enrichment of upregulated genes, located at Mb bin 118-120 (p-value 0.0554), as well as to an enrichment of upregulated genes on chr11 up, bin 10-12 and bin 61-63 (p-values 0.0022 and 0.0013, respectively).

To further investigate translocation-specific effects, we performed single-cell RNA-seq (scRNA-seq) analyses. As expected, we detected approximately 11% of CCND1-positive cells in our samples, which we considered to carry the induced translocation. Interestingly, they do not cluster separately from CCND1-negative cells, suggesting that there is a high level of similarity between the CCND1-positive and -negative populations, possibly combined with substantial heterogeneity within the CCND1-positive population (fig 4j). Nevertheless, when analyzing the distribution of upregulated genes per chromosome in CCND1-positive versus-negative cells, we observed a strong overrepresentation of genes on chr11 down (fig 4k, supp table 8), which was not observed for any other chromosome. These results highly suggest that the upregulation of these genes is linked to the presence of the translocation itself. Finally, we also observed a bias of upregulated genes in Mb bins 82-84 and bins 118-120 on chr11 down (respective p-values 0.0335 and 0.0594) and in Mb bins 64-66 (p-value 0.0287) on chr11 up. The latter two regions are in line with our previous hypothesis based on gene expression data analysis of MCL patients. In conclusion, both in MCL patients and in *in vitro* translocation models we detect major gene expression alterations in chr11 down, coinciding but not limited to the regions that get closer to the IGH enhancer in 3D space. Based on these results, we consider that this entire chromosomal segment, covering over 50 Mb, harbors a more permissive gene regulatory environment induced by the IGH enhancer. Interestingly, the large majority of the upregulated genes on this chromosomal segment in the single-cell assay already contain an active promoter in wildtype GM12878 cells (90.2%, 46/51 genes, supp table 8), which is strikingly more than expected by chance (46.8%, 214/457 genes), suggesting that the IGH enhancer has limited capacity to change promoter state - except within its neo-TAD leading to CCND1 overexpression - but rather enhances gene expression of active genes. Interestingly, all translocation-induced upregulated genes in Mb bin 64-66 on chr11 up, which is subject to a large change in radial positioning, also already displayed an active promoter prior to translocation formation (supp table 8). Altogether our results stress the importance of the epigenetic state of the cell in which this translocation occurs, both at the 3D genome and genome activation level, as it will determine its downstream gene expression effects and thus oncogenic transformation.

## Discussion

It is commonly known that SVs, such as translocations, can play a key role in disease development, particularly in malignancies^10,12,33^. However, the complex, genome-wide relationship between SVs, genome reorganization, and gene expression alterations in tumors remains largely unknown. In this work, we provide a detailed picture of the link between these layers in the context of MCL-associated translocations (fig 5). While chromosome conformation capture techniques, especially in combination with long-read sequencing, are utilized more and more to resolve complex karyotypes in cancer^34^, our results demonstrate that *in vitro* SV generation and downstream analysis are key to providing a holistic view of the effects of SVs on genome regulation. By using this approach, we show that MCL-associated IGH translocations upregulate a large set of genes on the entire affected chromosome arm downstream of the IGH locus. Thus, the full segment of chr11 down, spanning over 50 Mb, becomes influenced by the strong gene regulatory capacity of the IGH enhancer. So far, the only reported effects of IGH translocations *in cis* are those affecting single genes close to the breakpoint, namely CCND1 in MCL, and BCL6, BCL2 or MYC among others in other NHLs^18,35–37^. While our study focuses on understanding the very early effects of lymphoma formation, our results have a far more general implication, as they show for the first time that a strong enhancer can have a very broad long-range gene regulatory impact. Because SVs affecting strong enhancers are frequently found in cancer, we believe that our findings are widely applicable to understanding the impact of a large landscape of SVs beyond translocations, representing a myriad of tumor types. Furthermore, SVs are among the early events in tumor formation, revealing that their genome-wide effects are essential to further unveiling the precise mechanisms driving early carcinogenesis.

**Figure 5.**
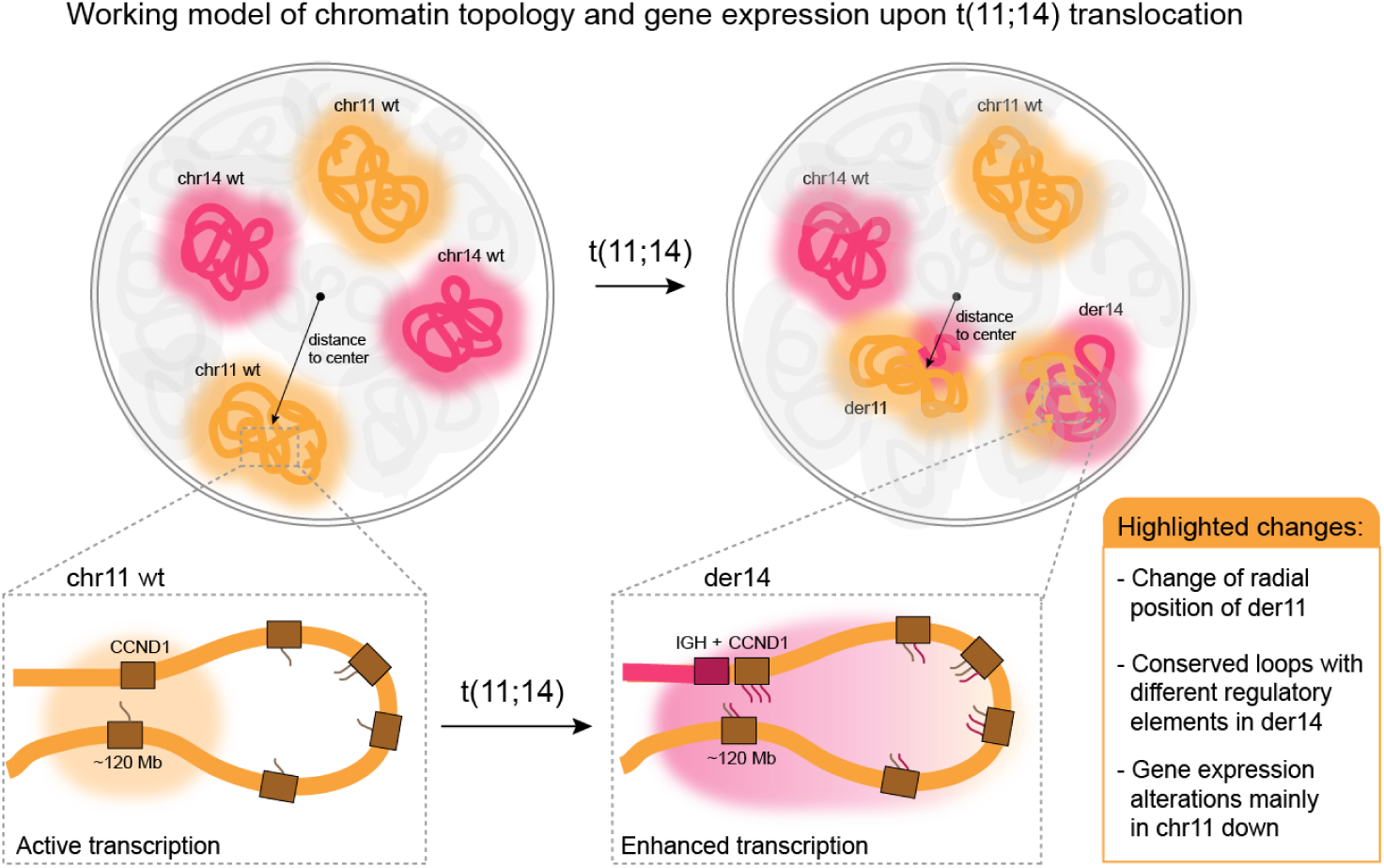
Working model. Both translocated chromosomes, derivative 11 (der11) and derivative 14 (der14), formed upon the t(11;14) in MCL undergo genome organization changes compared to wildtype, with der11 moving towards the nuclear center and der14 showing *de novo* intrachromosomal loop formation close and distant from the breakpoints. We furthermore show that the latter causes a pattern of enhanced transcription (indicated by the pink mRNA transcripts) affecting the entire segment of chromosome 11 downstream of the breakpoint, with the bins around Mb 120 being most susceptible to upregulate gene expression.

While the results from our genetically engineered cells unveiled that *de novo* translocations can lead to gene expression upregulation, with a major effect in derivative 14 downstream of the translocation breakpoint, not all genes on this chromosomal segment become upregulated. Instead, we show that pre-existing genome organization and activation prior to translocation formation are key for this downstream effect. For example, genes most susceptible to change expression are those located at Mb 118-120 on chr11 down, which are part of a conserved, long-range loop spanning 50 Mb that links these genes to the *CCND1* locus in 3D space. Upon translocation formation, this conserved loop leads to the establishment of long-range interactions between the strong IGH enhancer and the abovementioned cluster of genes. Interestingly, we show that the translocation-induced expression effects are not limited to this highly susceptible region, but can affect genes on the entire segment of chr11 down. Further research is necessary to explore whether all genes in this segment are simultaneously upregulated in individual cells or whether heterogeneity exists. Our single-cell data may point to the latter, although the intrinsic sparsity of this data does not allow us to draw definitive conclusions. Importantly, the induced gene expression effect is largely limited to genes with active promoters prior to translocation induction, highlighting that close proximity to the strong IGH enhancer alone cannot alter gene expression independently of a predetermined setup. *CCND1*, found next to the IGH enhancer within the neo-TAD, is an exception to this rule, being one of the few genes changing its promoter activity, which induces its expression upon translocation formation. Overall, based on our data, we speculate that the close proximity of the IGH enhancer drives gene expression enhancement in chr11 down. To the best of our knowledge, this represents a so far unreported phenomenon in which cancer cells can utilize translocations to hijack existing long-range interactions over tens of megabases and alter gene expression beyond changes close to the breakpoint.

We furthermore unveiled that MCL cells present an altered CT positioning pattern compared to healthy B cells (fig 5). In line with previous studies that widely report short and gene-rich chromosomes to occupy a more central nuclear position^8,38–40^, we detected that derivative 11, which drastically decreases size and increases gene density, moves towards the nuclear center. Simultaneously, we show that its regions close to the breakpoint display an enriched number of upregulated genes in MCL patient samples (Mb bin 67-69) and in our *in vitro* translocation models (Mb bin 64-66). Our results are in agreement with Bintou et al. who reported that der11 moves towards the nuclear center, and detected an upregulation of more than 80 genes in chromosome 11, in the GRANTA-519 MCL cell line and four MCL patient samples compared to a healthy LCL cell line (LCL RPMI8866). Similarly to chr11 up, all translocation-induced upregulated genes in Mb bin 64-66 harbored an active promoter prior to translocation formation.

Our findings have major consequences for understanding tumor formation, since the functional effects of translocations are likely dependent on pre-existing epigenetic characteristics to induce oncogenic effects. This finding is particularly important in the current era in which single-cell approaches enable the identification of more and more cell subtypes. Based on our findings, we may be able to predict translocation-induced effects in these subtypes, allowing us to better pinpoint tumoral cells of origin. In MCL cases, for example, translocations occur in early B cells in the bone marrow, but they do not fully transform into MCL cells until they reach the naive B cell stage. Therefore, our observations open doors to hypothesize that the genome activity state of naive B cells may provide key triggers to upregulate specific genes on chromosome 11 that drive tumor development. Notably, overexpressed genes in MCL patients are enriched in the exact same chromosomal regions as those genes affected by *in vitro*-induced translocations. This underlines the potential relevance of our findings for lymphomagenesis. Moreover, it provides a clear set of genes to be explored for their oncogenic effects and to design better murine MCL models, as well as therapeutic strategies in the context of MCL that is so far considered incurable. Upregulated genes in Mb bin 118-120 on chromosome 11, for example, emphasize the activation of T-cell pathways in MCL cells, including upregulation of CD3. While the expression of the T-cell marker CD5 is a common feature in MCL, cases with cytoplasmic expression of CD3 have been only sporadically reported^41,42^. Interestingly, T/B biphenotypic lymphocytes were recently revealed by Zhang *et al*.^43^ to be found in healthy murine and human blood samples. In the same study, these cells were described to originate from a subset of mature B cells and to express both CD5 and CD3 markers. CD5+/CD3+ cells furthermore have been reported to represent B-1a cells related to autoimmune disease^44^. Altogether, these results might indicate a link between T/B biphenotypic lymphocytes, CD5+/CD3+ B-1a cells and CD5+/CD3+ MCL cells. However, follow-up studies are needed to investigate whether these cells could be related to the cell of origin of MCL or how T-cell activation by other means could contribute to MCL pathogenesis.

In conclusion, the novel, broad-reaching changes of IGH translocations in genome organization, together with the associated transcriptional alterations emphasize the importance of genomic integrity to prevent disease development. Future research will reveal similar mechanisms by which other translocations or more broadly SVs influence chromatin structure, and uncover how these changes can contribute to tumor development. These findings will be crucial for developing better disease models, more precise (pre-)diagnostic tools, and new therapeutic strategies, due to a better global understanding of the early onset of malignant transformation.

## Author contributions

AO, RZ and KQ generated Z-138 Hi-C data. AO, MK and MO generated Tiled-C data. AO and NH generated 3D FISH data. RZ, CL and SB generated Z-138 allele-specific genotyping data. RZ and AR generated GM12878 *in vitro* translocations and gene expression data. AO, KQ, JRHM, FS, AV and RB analyzed MCL, CLL and normal B-cell Hi-C data. AO, HT and RB generated and/or analyzed 3D modeling data. AO and AS analyzed Tiled-C data. AO, EW, RG, NH and MB analyzed 3D FISH data. AO, AB and RB analyzed MCL patient RNA-seq data. AO, HT, JRHM and RB analyzed Z38 allele-specific Hi-C data. AO, AB, LA and RB analyzed *in vitro* translocation (sc)RNA-seq data. RB conceptualized the study, with contributions of AO and RZ. RB supervised the study. RB and AO wrote the manuscript. All co-authors read and commented on the manuscript.

## Conflicts of interest

The authors declare no conflicts of interest.

## Acknowledgments

The authors would like to thank Shyam Sundar Ramasamy (support on Tiled-C data generation), Jonas Paulsen (input on Chrom3D pipeline), Òscar Fornás and Eva Julià (support on chromosome sorting), Iana Kim (input on Tiled-C data analysis), Freddy Monteiro and the IRB Barcelona Functional Genomics Core Facility (support on Singleron scRNA-seq data generation), and Juan Valcárcel for advice on the manuscript. We acknowledge the support of the Spanish Ministry of Science and Innovation to the EMBL partnership, the Centro de Excelencia Severo Ochoa and the CERCA Programme/Generalitat de Catalunya. We acknowledge support from the Core Technologies Programme, the CRG Advanced Light Microscopy, Genomics and Bioinformatics Units, and the CRG/UPF Flow Cytometry Unit. This research was supported by grants from the European Research Council (ERC, LymphoTOP grant agreement number 101039265 to RB and RADIALIS grant agreement number 101088408 to MB). Views and opinions expressed are those of the authors only and do not necessarily reflect those of the European Union or the European Research Council Executive Agency. Neither the European Union nor the granting authority can be held responsible for them. This research was furthermore supported by grants from the Swedish Cancer Research Foundation (Cancerfonden, grant agreement number 22 2240 Pj 01 H to MB) and the Swedish Research Council (grant agreement number 2020-02657_3 to MB). RB was supported by a Junior Leader Fellowship from the la Caixa foundation. AO was supported by an EMBO scientific exchange grant (10156). AB is supported by an FPI fellowship from the Spanish Ministry of Science and Innovation (PRE2019-087574). LA is supported by a fellowship from the EMERALD International PhD Programme for MDs (101034290) and CEX2020-001049-S financed by MCIN/AEI/10.13039/501100011033.

## Supplementary Figure Legends

**Supplementary Figure 1.** The interchromosomal 3D landscape in MCL. **A)** Dissimilarity matrix comparing the Hi-C matrices of interchromosomal reads among samples, excluding reads between translocated segments (11down with 14up and 11up with 14 down). **B)** Log2 fold-change of the ratio of interaction frequencies between CLL and healthy B cells. Dashed lines highlight translocated chromosomes, again divided into “up” and “down” as in figure 1C. down = downstream of the breakpoint position, up = upstream of the breakpoint position, IF = interaction frequency.

**Supplementary Figure 2.** Allele-specific analysis of translocated chromosomes in MCL. **A)** Bivariate flow karyotype of Z-138 cells showing the isolated chromosome clouds. P2 and P3 represent wild-type chromosomes 14 and 11 respectively, while P1 indicates der11. **B)** Flowchart of SNP classification per chromosomal segment. SNP array, Hi-C and bivariate flow karyotyping sequencing data were used to identify heterozygous SNPs that were further used to genotype wild-type and translocated alleles based on bivariate flow karyotyping sequencing data. **C)** Fraction of Hi-C interactions in translocated versus wild-type alleles in Z-138 cells. Deviations from the dashed line (diagonal indicating the same fractions in wild-type and translocated chromosomes) represent allele-specific biases. **D)** Density plots of the relative distribution of radial position of chromosome 11 and 14 in JVM-2 compared to control (GM12878), 0.0 indicates the closest pixel to the lamina and 1.0 the most central position. Dashed lines represent the median. The positioning of chromosome 14 was assessed using the IGH break-apart probe, following the same principle as shown in figure 2D, but then for the IGH locus. **E)** Correlation of non-translocated chromosome positions among MCL, CLL and healthy samples. wt = wild-type, tx = translocated, down = downstream of the breakpoint position, up = upstream of the breakpoint position, ns = not significant.

**Supplementary Figure 3.** Intrachromosomal interaction landscape of translocated chromosomes in MCL. **A)** PCA of triplicates used in Tiled-C analysis, based on their intrachromosomal interactions within the targeted region. **B)** Tiled-C read count heatmaps in chromosome 11 comparing the U-266 cell line (right-upper triangle) with GM12878 control cells (left-bottom triangle). Dashed lines highlight the region where the IGH enhancer is inserted in chromosome 11 in U-266. Arrows indicate differences in contact strength from the IGH enhancer insertion to chromosome 11. **C-D)** Same as in B, but depicting derivatives 14 and 18 comparing KARPAS422 and Pfeiffer cell lines carrying the t(14;18) translocation (right-upper triangle) with the control GM12878 (left-bottom triangle). **E-G)** Differences of insulation score on chromosomes chr14 up and chr18 down in the analyzed cell lines compared to GM12878 cells. Heatmaps represent interaction frequencies from Tiled-C read counts in the depicted areas, shown with a red square in the chromosome representation on top. Colors from the line plots indicate a higher (blue) or lower (red) insulation score. der = derivative, wt = wild-type, tx = translocated, down = downstream of the breakpoint position, up = upstream of the breakpoint position, IS = insulation score, ZJ = Z-138 and JVM-2, KP = KARPAS422 and Pfeiffer.

**Supplementary Figure 4.** Gene expression analysis in cells carrying the t(11;14) translocation. **A)** Graphic representation of the location of up- and downregulated genes in MCL patients throughout chr4 up. Names for upregulated genes with a log2FC fold change higher than 1 are depicted. Below, the compartment score of each Mb bin is represented, whereby blue represents B compartments, and red A compartments. **B)** Proportion of genes from the dataset that are associated with upregulated pathways by GO analysis. **C)** Network plot, nodes represent the genes linked to the enriched pathways depicted in B, and edges indicate the genes that are connected through the same pathways. **D)** Graphical representation (left) and digital PCR results (right) indicating the estimated percentage of translocated cells after *in vitro* translocation generation in each replicate. **E)** Level of CCND1 upregulation after *in vitro* translocation generation compared to different controls. **F)** VSD-transformed gene expression levels of significantly variable genes in translocated replicates (Tx1-Tx3) compared to controls 1-6 (C1-C6) (FDR < 0.1, FC > 1.1). **G)** Fraction of upregulated genes per chromosome in translocated samples compared to C1-C6 controls (FDR < 0.1, FC > 1.1). Dashed line represents the expected fraction of genes. down = downstream of the breakpoint position, up = upstream of the breakpoint position.

## Supplementary Table Legends

**Supplementary Table 1. Sample characteristics.** Characteristics of the primary samples used in the study as well as the reference to previous studies in which the samples were used. MCL = mantle cell lymphoma, CLL = chronic lymphocytic leukemia, NA = non-assigned.

**Supplementary Table 2. Structural variants in MCL samples.** List of patient-specific structural variants found in the MCL patients included in this study, as defined by Nadeu *et al*, Blood 2020 (WGS) and Beà *et al*, PNAS 2013 (WGS, WES, SNP arrays).

**Supplementary Table 3. Common altered interchromosomal interaction in MCL.** List of common top 10% of interchromosomal interactions altered in MCL versus healthy B cells and CLL.

**Supplementary Table 4. Heterozygous SNPs in MCL cell line Z-138.** Full list of heterozygous SNPs on chromosomal segments chr11 up, chr11 down and chr14 up, including genotype information on the non-translocated and translocated alleles.

**Supplementary Table 5. Number of allele-specific reads on chromosomes 11 and 14 in Z-138.** Number of Hi-C reads in Z-138 that map to either the non-translocated or translocated chromosome 11 or 14, representing an interchromosomal ligation product.

**Supplementary Table 6. Differentially expressed genes in MCL samples compared to healthy B cells.** Differentially expressed protein-coding genes in MCL compared to mature healthy B cells, log2FC > 1, FDR < 0.1. Upregulated genes are expressed in at least 3 MCL patients.

**Supplementary Table 7. Differentially expressed genes upon *in vitro* translocation generation bulk.** Differentially expressed protein-coding genes in GM12878 populations carrying ∼10% translocated cells versus control GM12878 cells, log2FC > 1.1, FDR < 0.1.

**Supplementary Table 8. Differentially expressed genes upon *in vitro* translocation generation single-cell.** Differentially expressed protein-coding genes in CCND1-positive GM12878 cells upon *in vitro* translocation generation, compared to CCND1-negative cells using scRNA-seq, FDR < 0.1.

**Supplementary Table 9. IGH translocation breakpoints in cell lines.** IGH translocation breakpoints or IGH insertion locations in Z-138, JVM-2, U-266, KARPAS422 and Pfeiffer cell lines, defined using the Tiled-C data at 5 kb resolution.

**Supplementary Table 10. Break-apart probes.** Characteristics of break-apart probes targeting the *CCND1* and IGH loci for FISH experiments.

## Methods

### 1. Samples and cell lines

#### Primary samples

No primary samples were processed for experiments in this study. All Hi-C and RNA-seq data of patient samples and healthy donors were mined from publicly available data from a previous publication^24^. Sample characteristics in supplementary table 1.

#### Cell lines

Cell lines Z-138, JVM2 and Pfeiffer were obtained from ATCC (cat. no. CRL-3001; cat. no. CLR-3002; cat. no. CRL-2632), U266 from DSMZ (cat. no. ACC 9), KARPAS422 from Merck (cat. no. CB_06101702), and GM12878 from Coriell (cat. no. CVCL_7526). Z-138 cells were cultured in Iscove’s Modified Dulbecco’s Medium containing 4 mM L-glutamine (Gibco^TM^, cat. no. 31980030), supplemented with 10% horse serum (Gibo^TM^, cat. no. 10368902) and 1% penicillin/streptomycin (Gibco^TM^, cat. no. 15140122). JVM2, U266 and GM12878 cells, were cultured in RPMI-1640 Medium 2 mM L-glutamine (Gibco^TM^, cat. no. 21875034) containing 15% fetal bovine serum (FBS) (Gibco^TM^, cat. no. 10270106) and 1% penicillin/streptomycin (Gibco^TM^, cat. no. 15140122). KARPAS422 and Pfeiffer cells were cultured in RPMI-1640 medium with 20% and 10% FBS respectively. All cells were cultured at 37°C under 5% CO_2_ and atmospheric O_2_ conditions, and they were regularly passed and tested for mycoplasma infection.

### 2. Bivariate flow karyotyping, chromosome sorting and sequencing

Z-138 cells were prepared for flow karyotyping as described^45^ with few modifications. Briefly, 20 million cells were treated with 0.1 μg/ml Colcemid (Sigma-Aldrich, cat. no. 4725301) for 6 h at 37°C. Pelleted cells were resuspended in 10 ml hypotonic solution (75 mM KCl, 10 mM MgSO4, 0.2 nM spermine and 0.5 nM spermidine, pH 8.0) and incubated for 27 min at 37°C. After centrifugation (5 min at 300 g), the cell pellet was resuspended in 1 ml cold polyamine solution buffer (15 mM Tris pH 8.0, 2 mM EDTA, 0.5 mM EGTA, 80 mM KCl, 3 mM DTT, 0.25% Triton X-100, 0.2 mM spermine and 0.5 mM spermidine, pH 8.0) and incubated on ice for 35 min to release the chromosomes. Next, samples were vigorously vortexed for 30 s and filtered through a 35 μm mesh filter (Falcon, cat. no. 352235). Chromosomes were double stained by adding 10 μl of 1M MgSO_4_.7H_2_O, 40 μg/ml of chromomycin A3 (Sigma-Aldrich, cat. no. C2659), an AT-bp specific dye, and 5 μg/ml of Hoechst 33258 (Sigma-Aldrich, cat. no. H3569), a CG-bp specific dye, for 8 h at 4°C. Before sorting, samples were incubated with 10 mM of sodium citrate and 25 mM of sodium sulfite for 2 h on ice. Flow karyotyping for chromosome sorting was performed on the BD Influx cell sorter (Becton Dickinson, San Jose, CA) using the UV (355 nm) and Deep Blue (457 nm) high laser power and 100 μm nozzle. Sorted chromosomes were treated with 200 μg/ml of RNAse (NEB, cat. no. EN0531) for 30 min at 37°C and 200 μg/ml with Proteinase K (Invitrogen, cat. no. AM2546) for 16 h at 50°C. After treatment, the buffer was exchanged by dialysis using Pur-A-Lyzer™ Maxi Dialysis Kit (Sigma-Aldrich, cat. no. PURX50005) against 1x TE buffer for 48 h at RT. To reduce the volume, samples were concentrated by evaporation in a SpeedVac Vacuum concentrator up to 100 μl volume and purified with 2x SPRI beads (Beckman Coulter™ Agencourt AMPure XP, cat. no. A63880). Sequencing libraries were prepared using the NEBNext Ultra II FS DNA Library Prep Kit (NEB, cat. no. E7805). Libraries of around 300 bp size were sequenced using the Illumina HiSeq 2500 system (paired-end, 2x 125 bp). Libraries corresponding to the wildtype chromosomes 9 to 12 and der11 were sequenced for a total of 133 and 43.4 million reads, respectively.

### 3. Hi-C data generation

Hi-C was performed on Z-138 as previously described^46^ with minor modifications reported here. Briefly, 30 million cells were cross-linked with 1% formaldehyde (Merck, cat. no. F8775-4X25ML), quenched with 0.125 M glycine solution (Sigma-Aldrich, cat. no. 50046) and flash-frozen in dry ice. Samples were then stored at −80°C until further steps were performed. After lysis, chromatin was digested using two restriction enzyme conditions, with two or three biological replicates per condition. Cells were either digested with 72 µl MboI (25 U/µl, NEB, cat. no. R0147M), or a mix of three restriction enzymes: 30 µl of NcoI-HF (20 U/µl, NEB, cat. no. R3193S), 30 µl of BclI-HF (20 U/µl, NEB, cat. no. R3160S) and 30 µl BsrGI-HF (20 U/µl, NEB, cat. no. R3575S) and incubated overnight at 37°C without shaking. All the enzymes were added at two different time-points to refresh their enzymatic activity, except for BsrGI, which was only added once, as its enzymatic activity lasts longer than 8 h. Next, enzymes were heat-inactivated for 20 min at 65°C (MboI) or 80°C (combination of three restriction enzymes). Finally, chromatin was extracted using the ethanol extraction method and resuspended in a final volume of 400 µl. The final concentration was calculated by Qubit (dsDNA High Sensitivity Assay) and ligation and digestion efficiency were checked on a 0.6% agarose gel.

Hi-C libraries were built from 4 µg of DNA per sample and sonicated using a BioruptorPico device (Diagenode, cat. no. B01060010, 3 cycles, 20 s on, and 90 s off). The efficiency of sonication was checked on a 1.2% agarose gel for 20 min. Sheared samples were cleaned up using 1/3x pre-washed Streptavidin T1 beads (T1 Dynabeads MyOne, Invitrogen, cat. no. 65601), incubated for 30 min in 100 µl of the end-repair library mix (10 µl of 1x T4 DNA ligase buffer with 10 mM ATP (10x, NEB, cat. no. B0202S), 2 µl of dNTPs (25 mM each), 5 µl of T4 PNK (10,000 U/ml, NEB, cat. no. M0201S), 4 µl of T4 DNA polymerase (3,000 U/ml, NEB, cat. no. M0203S) and 1 µl of the DNA polymerase I large (Klenow) fragment (NEB, cat. no. M0210L)), and reclaimed with 100 µl of A-tailing mix (10 µl of 10x NEBuffer2; 5 µl of 10 mM dATP; and 5 µl of the Exo minus Klenow fragment (3’ 5’), 5,000 U/ml, NEB, cat. no. M0212S) for 30 min at 37°C without shaking. Illumina loop adaptors (2.5 µl, NEBNext Multiplex Oligos for Illumina, Index primer set 1, NEB, cat. no. E7335S) were ligated to the DNA fragments in 50 µl of the Quick ligation mix containing 25 µl of 2x Quick Ligase reaction buffer (NEB, cat. no. B2200S) and 2 µl of T4 DNA ligase (2,000,000 U/ml, NEB, cat. no. M2200M), for 15 min at RT with 3 µl of the USER adaptor (NEBNext Multiplex Oligos for Illumina, NEB, cat. no. E7338A) for an additional 15 min at 37°C. Final libraries were resuspended in 10 µl of EB buffer (10 mM Tris-Cl, pH 8) from Qiagen (cat. no. 19086) and amplified using 2.5 µl of Universal PCR primer (NEBNext Multiplex Oligos for Illumina, NEB, cat. no. E6861A), 2.5 µl of indexed primers (NEBNext Multiplex Oligos for Illumina, Index primer set 1, NEB, cat. no. E7335), and the NEBNext High Fidelity PCR Master Mix (12.5 µl of 2x master mix, NEB, cat. no. M0541S), during 8 PCR cycles (10 s of denaturalization at 98°C, 30 s of annealing at 65°C and 30 s of elongation at 72°C). The final libraries were extracted using 70% EtOH and resuspended in 30 µl EB buffer, quantified by Qubit (dsDNA High Sensitivity Assay) and profiles were evaluated on the Bioanalyzer 2100. Libraries were then sequenced on a HiSeq2500 (50 bp, paired-end reads, 60 million reads per library on average).

### 4. Hi-C and bivariate flow karyotype data processing and allele-specific Hi-C analysis

#### Hi-C data processing

Hi-C data of 5 individual MCLs, 7 individual CLLs, 12 individual normal B-cells, and 5 Z-138 replicates were processed. Hi-C reads were iteratively mapped using TADbit mapper^47^ against genome build GRCh38. Mapped reads were parsed and duplicated reads or reads with a distance of less than 1 Kbp between the two read pairs were filtered out. Valid pairs were then normalized using TADbit norm at two different resolutions, 100 Kb and 1 Mb (’Vanilla’ normalization, >1,000 reads per bin). Biases files (created by the TADbit norm command) and bam files with valid pairs (created by the TADbit mapper command) were used for further processing. First, after obtaining raw contact frequency matrices per individual primary sample using the TADbit bin function at 1 Mb resolution, these were normalized to a total interaction count of 150 M per sample. Next, these matrices were merged (using the median interaction frequency per interaction bin) to obtain three merged raw contact matrices, one for MCL (n=5), one for CLL (n=7), and one for normal B cells (n=12). Furthermore, compartments were assigned per sample using the TADbit segment function (using the 100 Kbp resolution data). Per chromosome and per sample, the eigenvectors representing the compartment scores were defined by selecting the one with a high correlation (∼0.6 or higher) between the eigenvalues and GC content. In case the correlation with GC content was similar for multiple eigenvectors and/or when the difference in the percent of variation explained by different eigenvectors was less than 1.5 fold, visual inspection was conducted on Pearson’s heatmaps of the eigenvectors. To obtain one compartment score per 1 Mbp bin in each individual sample, the median compartment scores of ten 100 Kbp bins were defined. Finally, to obtain one compartment score per sample type (MCL, CLL or normal B cells), the median compartment scores were calculated to merge the individual 1 Mbp scores. A and B compartments were assigned having a positive or negative eigenvalue respectively.

#### Interchromosomal interaction frequencies in primary samples

First of all, to avoid artifacts from regions with low-mappability, interactions from outlier 1 Mb bins with < 60.000 interactions in total were flagged to be ignored (set to NA - non-applicable). These mainly represent telomeric and centromeric regions. Next, to obtain pairwise interchromosomal interaction frequencies between chromosomes in MCL, CLL and normal B cells, we calculated the total interaction counts of the merged matrices per chromosome pair and divided this number by the product of the total number of Mb bins per chromosome (scaling the interaction frequency per square Mbp bin). Log2FC of the interaction frequencies was calculated by dividing the obtained values in MCL by the values obtained in MCL or CLL.

#### Bivariate flow karyotype data analysis Z-138

Reads of the sorted chromosomes were trimmed using TrimGalore (version 0.6.6), with a minimal Phred quality score of 13, a maximal allowed error rate of 0.1, a stringency of 3 and a minimal read length of 35 bp. Quality checks of the reads before and after trimming were carried out by fastQC (version 0.11.9). Next, trimmed reads were aligned against the reference GRCh38 human genome using hisat2 (version 2.2.1), with default parameters, a minimal allowed intron length of 20 bp and a maximal of 500,000 bp. Then, genotype calling was carried out following the variant discovery workflow present in the GATK Best Practices recommendations (https://gatk.broadinstitute.org/hc/en-us/sections/360007226651-Best-Practices-Workflows). After marking duplicates (’MarkDuplicatesSpark’), the base quality scores were recalibrated first by producing the recalibration table (’BaseRecalibrator’ using the known polymorphic sites from the GRCh38 104 human genome release version) and then by applying the recalibration (’ApplyBQSR’) to the original BAM files with the aligned reads. Variants were then called independently for each chromosome sorting sample, limiting the calling to chromosomes 11 and 14, and using ‘HaplotypeCaller’ in GVCF mode. Next, these per-sample variants were joined creating a database with all the consolidated per-sample variants (’GenomicsDBImport’) and the joined genotyping with ‘GenotypeGVCFs’. All these arguments were launched with default arguments. The used GATK version was 4.2.0.0, and its associated Picard version was 2.25.0. Based on the bivariate flow karyotype data we also assigned the chromosomal breakpoint regions in Z-138, resulting in the following assignments in this cell line: chr11 up <69,582,000, chr11 down >69,583,000, chr14 up <105,890,000, 14 down >105,891,000 Mb.

#### Heterozygous SNP assignments and allele-specific genotype calls in Z-138

Heterozygous SNPs on chromosomes 11 and 14 in Z-138 were determined using a combination of SNP array, bivariate flow karyotype, and Hi-C data. The latter two were generated in this study, while the former data was produced in the context of a previous publication^28^. The SNP array data of two replicates of Z-138 was analyzed using the *crlmm* R package (v. 1.58.0). Default quality scores for SNPs (>0.7) and samples (>5) were used as filtering thresholds. SNPs assigned as heterozygous in both replicates were considered further. Similarly to the processing of the fastQ files of the bivariate flow karyotype data described above, Hi-C data was processed to obtain heterozygous SNP calls. Bam files of the 5 replicates were merged after base recalibration followed by variant calling in GVCF mode and selection of heterozygous SNPs with a quality score above the default threshold in GATK. Only for chr11 up, heterozygous SNPs could also be determined using the bivariate flow karyotype data. To that end, genetic variants with a genotype quality score equal to or greater than 20 were used to retrieve the genotype information of the wildtype chromosome 11 and the der11. By comparing the obtained genotypes, positions with heterozygous SNPs were defined. Next, for each heterozygous SNP on chromosomes 11 and 14, the genetic variants with a genotype quality score equal to or greater than 20 of the chromosome sorting data were used to assign a specific genotype call to derivatives 11 and 14 (supp fig 2B, supp table 4).

#### Allele-specific Hi-C analysis Z-138

The Hi-C read sequences covering heterozygous SNP with allele-specific genotype calls were consequently extracted from the merged bam file, containing the reads of all 5 Z-138 Hi-C replicates. Using Bedtools (version v2.27.1) intersect, the BAM files were screened for overlaps with the heterozygous SNP positions, whereby only reads with at least one overlap were kept. Next, the base calling of the heterozygous SNP position was retrieved from the read sequences. Alleles were assigned to the wild-type or derivative chromosome by comparing the SNP base pair of the read with the assigned reference and alternate base calls. Finally, the number of interactions from the wild-type and translocated fragments of chromosome 14 and 11 to each chromosome was calculated.

### 5. Tiled-C data generation and analysis

#### Viewpoint selection and oligo pool preparation

To determine the regions of interest, we evaluated publicly available Hi-C data of GM12878 cells from Rao *et al*.^4^. We selected regions comprising 5 TADs surrounding the breakpoints in chromosomes 11, 14 and 18, and TAD borders were further evaluated using another publicly available dataset^48^. Then, oligo pools targeting DpnII cutting sites within the selected regions (hg38: chr11: 66,940,000 - 71,860,000; chr14: 103,980,000 - 107,043,718; chr18: 61,460,000 - 64,500,000) were designed using the Oligo tool^31^ and filtered for repetitive sequences. Double-stranded, biotinylated DNA oligonucloetides were purchased from Twist Biosciences (Twist NGS Target Enrichment Oligonucleotide Panel, cat. No. 100533), and reconstituted following manufacturer’s instructions to a stock concentration of ≥ 1 µM for each unique oligonucleotide.

#### Sample preparation

Tiled-C was performed as previously described^32^ with minor modifications. Briefly, 5 million cells were collected in single-cell suspensions, fixed in 2% (vol/vol) formaldehyde and incubated for 10 min at RT while tumbling or rotating. Formaldehyde was quenched with 1 M cold glycine, and the cell pellet was lysed in ice-cold lysis buffer (1 M Tris, pH 8; 5 M NaCl; 10% (vol/vol) Igepal CA-630; 1 tablet of cOmplete EDTA-free Protease Inhibitor Cocktail tablets, Roche, cat. no. 04693132001) for 30 min. Cells were pelleted and resuspended again in 1x DpnII buffer (NEB, cat. no. B0543), snap-frozen and stored at −80℃ for short-term storage. Digestion with DpnII enzyme (NEB, cat. no. R0543) was performed as described^32^, followed by ligation using the NEBNext Quick Ligation kit (NEB, E6065) and DNA extraction with phenol-chloroform-isoamyl alcohol (PCI) and ethanol precipitation. DNA pellets were resuspended in TE buffer and kept at −20℃ until library preparation. Quality control of each step was performed in a 15 (wt/vol) agarose gel using 1x Tris acetate-EDTA buffer, run at a moderate speed (70 mA) using 1 μl GeneRuler DNA ladder (Thermo Scientific, cat. no. SM0331), 15 μl of each control and 5 μl of 3C library. Qubit dsDNA High-Sensitivity assay kit (Thermo Scientific, cat. no. Q32854) was used to determine the final DNA concentration, and a standard yield of 60-75% of input DNA was obtained (> 15 μg from 5 million cells). Only libraries with >70% digestion efficiency were taken for the next step.

#### Library preparation and sequencing

Before library preparation, 6.5 μg of DNA for each sample was diluted in 1x TE buffer and sonicated using Covaris to a final mean size of around 250 bp. 2 μl of the sample pre- and post-bead clean-up (SPRIselect beads, Beckman Coulter, cat. no. B23317) were run in the TapeStation D1000 as a quality control to check peak size. Then, samples were end-repaired and adaptor-ligated using the NEBNext Ultra DNA Library Prep Kit for Illumina (NEB, cat. no. E7645S), and indexed in duplicate by PCR using the Herculase II Fusion DNA Polymerases kit (Agilent, cat. no. 600677) and the NEBNext Multiplex Oligos for Illumina, Primers Set 1 and 2 (NEB, cat. no. E7335 and E7500). We proceeded by pulling the samples together, and 2 μg of DNA from each sample were dried out using a Vacuum Speedback machine. Samples were then captured with the probe pools using the fast hybridization mix, fast hybridization enhancers and amplification primers from the Twist Fast Hybridization and Wash kit (Twist, cat. no. 104180), and the blocker solution and universal blocker from the Twist Universal Blockers kit (Twist, cat. no. 100578). After streptavidin washes using MyOne Streptavidin C1 Dynabeads (ThermoFisher, cat. no. 65001), capture amplification was performed by PCR of 22.5 μl per sample in 9 cycles, using the KAPA HiFi HotStart ReadyMix kit (Roche, cat. no. KK2602). After AMPure XP beads clean-up, DNA was purified with ethanol and resuspended in PCR-grade water. Final library quantification was performed using high-sensitivity Qubit assay and the KAPA Library Quantification Kit (Roche, cat. no. KK4824) with 1:10,000 and 1:20,000 dilutions. Fragment analyzer to detect the size of the final peaks was also performed. Libraries were diluted to a final concentration of 4 nM and sequenced with the NextSeq 500/550 HighOutput Kit v2.5 (75 cycles, Illumina, cat. no. 20024906), with 75 bp reads and pair-end sequencing, obtaining around 1 billion sequenced raw reads (62 million reads per sample in average).

#### Data analysis

Sequencing results were analyzed using the CapCruncher pipeline, as specified by Downes *et al*.^32^. Shortly, sequencing replicates were merged before alignment, and after a quality control test of the fastQ files, sequencing data was aligned to a custom hg38 genome containing an additional chromosome (named chromosome 23) constructed by concatenation of the targeted regions in chromosome 14, 11 and 18 respectively, and excluding alternative haplotypes. The captured regions were consequently masked from the original chromosomes to avoid multiple alignment errors. Following alignment, data was filtered for duplicates and unmapped reads using *pairtools*, and matrices from biological replicates were aggregated and scaled based on the number of restriction sites per fragment. Final read counts were normalized by iterative correction and eigenvector decomposition (ICE)^49^ using HiCExplorer (v. 3.7.2) and cooler (v. 0.9.3) packages. Next, breakpoint regions were defined based on the drop in contact frequency from one side of the translocation to the other (supp table 9). From the constructed chromosome 23, reads from each derivative or chromosomal fragment were selected and re-scaled to be able to compare them amongst cell lines. Tiled-C heatmaps were visualized using the HiContacts package (v. 1.6.0) in RStudio. From the output data, insulation scores (IS) on strong boundaries were calculated using Cooltools too (v. 0.6.1) and represented in R. TAD borders were called using HiCExplorer (v. 3.7.2).

### 6. FISH data generation

#### Sample preparation

Suspension cells were prepared as recommended by the probes’ manufacturer (Cytocell Ltd., Sysmex Group Company), with minor modifications to what was previously described (Nederlog, P.M. *et al*, 1989). Briefly, 1 million Z-138, JVM-2 and GM12878 cells were harvested and washed with 1x PBS. Then, 250 µl of freshly prepared cold fixative (3:1 methanol: glacial acetic acid) was added, and cells were incubated for 10 min at RT. Colcemid and hypotonic (KCl) treatments were avoided to maintain the interphase 3D nuclear structure as preserved as possible. Then, the supernatant was removed by centrifugation for 5 min at 1500 rpm, and 5 ml of cold fixative was added in a drop-wise manner while vortexing the sample. Cells were centrifuged again, and fixation was repeated until the cell pellet was white (2-3 times). Next, the cell pellet was suspended in fixative solution, mixed uniformly and spotted directly onto chambered coverglass (Nunc^TM^ Lab-Tek^TM^ II, Thermo Fisher, cat. no. 154526). The slide was left to air dry, and cell dispersion was assessed through a cell culture inverted microscope (DMi1, Leica Microsystems). Then, the slide was immersed in 2x SSC for 2 min at 37°C in a water bath, and directly placed in freshly-made pepsin working solution (20 µl of 10 mg/µl Pepsin, Roche, Sigma-Aldrich, cat. no. 10108057001, in 48 ml of distilled water (Ecolav, B. Braun, cat. no. 387874) containing 1 ml of 1 N HCl) for 40 s to 1 min. Slides were then washed with 1x PBS for 5 min and incubated with RNAse A (Thermo Fisher, cat. no. EN0531) diluted in 2x SSC (2 µl of RNAse A in 50 µl of 2x SSC) for 1 h at 37°C. Subsequently, the slides were washed with 2x SSC for 2 min at RT, and dehydrated successively in 70%, 85% and 100% ethanol dilutions. Next, slides were air-dried and 10 µl of fluorescent-probe mixtures pre-warmed for 5 min at 37°C were spotted onto the slide (see section below for probe details). The fixed cells were then covered with a glass coverslip (24×24mm, Menzel-Gläser, VWR, cat. no. 630-2104) and sealed with rubber solution glue (Marabu GmbH & CO, cat. no. 2901 10 000). DNA and probes were denaturalized together on a hotplate for 5 min at 78°C, with a weight on the slide, and covered from the light. Slides were then incubated overnight at 37°C in a humid, light-proof chamber inside an incubator. On the following day, coverslips and all traces of glue were removed, and slides were immersed in 0.4x SSC solution for 2 min at 72°C in a water bath. After washing the slides with 2x SSC + 0.05% Tween20 at RT for 30 s, slides were washed in 2x SSC for 5 min and allowed to air-dry. After that, 15 µl of antifade mounting media (Vectashield, Palex Medical SA, cat. no. 416397) were added to each slide, and a coverslip was applied and sealed with nail polish. Slides were directly imaged using a Zeiss LSM 980 microscope with airyscan, or stored at −20°C for short-term storage.

#### Probes

Break-apart probes for the *CCND1* and IGH loci were purchased from Cytocell (Sysmex Company Group, Oxford Gene Technology, cat. no. LPS 030 and LPH 014, respectively), which target upstream and downstream of the gene of interest with two fluorophores (supp table 10). For each experiment, 10 μl of the corresponding break-apart probes were mixed and pre-warmed for 5 min at 37°C. Then, the probe mixture was spotted onto the cell layer on each slide, and the following steps were performed as stated in the previous section.

#### Data acquisition

High-throughput imaging was performed using a fully motorized inverted Zeiss linescan confocal LSM980 unit with Airyscan 2 system, controlled by Zen Blue 3.2 software. Images were acquired using a Plan-Apochromat 63x/1.4 oil immersion objective and Zeiss 1.518 refractive index (Abbe number: 45) oil immersion media. For excitation, 405 nm (for DAPI), 488 nm (Alexa Fluor 488) and 561 nm (Texas Red) laser diodes were used. The emitted light was detected in a sequential mode with the Airyscan detector with the emission filters: 465 (DAPI), 517 (Alexa Fluor 488) and 592 (Texas Red) with an Airyscan detector. Images covering a 20 μm axial range completely covering the cell nuclei volume were acquired, with stacks of 0.13 μm and a voxel size of 73 x 73 x 130 nm^3^. Zoom factor was 3.6, scanning was unidirectional and line average was set to 2, with an image size of 35.7 x 35.7 μm^2^. Imaging was conducted in a controlled environment to maintain sample integrity, and exposure time that was maintained for all samples. Before acquiring images, 200 nm multi-color beads (Invitrogen™ TetraSpeck fluorescent microspheres, cat. no. T14792) were imaged with exactly the same settings and in the same environmental conditions, to correct for chromatic shift in the biological sample.

### 7. FISH analysis

#### Chromatic aberration correction

First, chromatic aberration was corrected closely following the procedure by Kozubek *et al*.^50^, using the TetraSpeck beads images which allowed for the detection of chromatic shifts between channels. To create a per-dataset correction, first, dots were identified in each channel of the bead images using a Difference of Gaussians (DoG) filter; second, dots were localized to sub-pixel precision by Maximum Likelihood (ML) fitting using a Gaussian model for the dot shape. Clustering was applied to find correspondent pairs, giving the coordinates of the same bead in two channels at a time. With correspondent pairs as input, a correction transform was found by the least squares method, specifically a polynomial correction of order 2 was used in the lateral plane and of order 0 (a constant shift) in the axial direction, which aligns all channels to DAPI. Then, the per-dataset correction transform was eventually applied to the coordinates of the detected dots, i.e. after the dot detection procedure. After correction, the largest mean square error between the correspondent pairs was found to be 0.25 pixels, indicating that subpixel precision was reached.

#### Nuclei segmentation

The DAPI channel was binarized using the following steps: (1) Otsu’s method was used to create an initial binarization in MATLAB (*imbinarize*, MathWorks, Natick, MA). (2) Then, Iterated Conditional Modes (ICM) were used to improve the initial segmentation^51^, and the mean variance of pixels classified as foreground with respect to the background from the previous step was used for initialization. The homogeneity parameter, β, was set to 1.5. (3) Face-connected regions were identified and their holes were filled using morphological operators. (4) Nuclei overlapping the image edges were discarded. (5) Nuclei were split using the watershed cuts (WSC) from Cousty *et al*.^52^. (6) The resulting masks were relabeled again. Finally, G1 cells were selected by excluding those cells in which the total DAPI intensity was outside the expected range (1e9-3.5e9), which was set manually by inspecting the histogram of the per-nuclei DAPI values.

#### Dot detection

Images were convolved with a Laplacian of Gaussian (LoG) filter with *σ = [1.55,1.55,3.29],* considering [σ_x_,σ_y_,σ_z_]. The image was then partitioned into patches by the watershed cuts (WSC) algorithm as explained previously^52^. For each patch, *P*, two features were used, *F1 = (max(P) - min(P))* and *F2 = - min(LoG(P))*, to separate the patches representing the true dots from the background. For each remaining patch, the center of mass (COM) was calculated. For this purpose, the min value was subtracted from the region, meaning that only the values higher than the min value of the local patch were used for the COM calculation. To avoid over-segmentation, known to possibly occur for probes covering large genomic regions, patches closer than *d_0_ = 12* pixels were merged.

#### Quality control

A final quality control was performed, and only those nuclei that met all criteria for each nucleus were kept for further analysis. First, each nucleus needs to have two pairs of dots, corresponding to the upstream and downstream segments of *CCND1*/IGH loci in the two alleles of a diploid cell line. Second, each cell needs to have two wild-type copies (GM12878) or a wild-type and a translocated copy (Z-138 and JVM-2). Due to the nature of the break-apart probes, dots will localize close together on a wild-type chromosome, while being further away when hybridizing to translocated chromosomes. Given that each of our cells carries at least one wild-type allele, we assigned the dot pair with minimum distance first, while the remaining dots formed the second pair. To properly assess whether the assigned dot pairs represent a wild-type (non-broken) or translocated (broken) chromosome, we built a model for the expected distance in the known wild-type case of GM12878 cells. Therefore, we estimated the chi-square-3 probability of a dot pair being wild-type using its square Euclidean distance, normalized for the σ_x_, σ_y_, and σ_z_ values estimated from GM12878. We took a threshold of 0.95 to reject the null hypothesis that pairs come from the non-broken wild-type chromosome, and using this threshold, we classified each dot pair as wild-type or translocated for all cells (2 dot pairs per cell). Wild-type nuclei are required to have two non-broken dot pairs, while translocated cells are required to have one non-broken and one broken dot pair.

#### Measurements

For each nucleus, we measured the distances between each dot and the closest region of the nuclear lamina (inferred from the DAPI signal). The Euclidean distance transform was applied as previously described^53^, taking the anisotropic pixel size into account. The distance values were scaled to have a maximum value of 1, meaning that all distance values are relative, where 0 means that the detected dot is located at the nuclear lamina, and 1 means located at the nuclear center. Finally, the distance maps were interpolated over the fitted coordinates of the detected FISH probes, and downstream analyses were performed in RStudio (version 2024.04.0+735).

### 8. Chrom3D modeling

#### Deconvolution of Hi-C matrices

To feed the modeling software with diploid models, the Hi-C matrices were deconvoluted as explained next. For the normal B cell and CLL merged (1 Mb resolution) Hi-C matrices, intrachromosomal reads were divided by two and assigned to what we call alleles a and b. Interchromosomal reads were divided by four and assigned to the four possible combinations of alleles (a-a, b-b, a-b, b-a). In MCL, for chromosomes 1-10, 12-13 and 15-22 the same approach was applied as described above. For chromosomes 11 and 14 in MCL, we assigned the same amount of reads to the single wildtype chromosome as observed in normal B cells. The remaining reads were assigned to the derivative alleles. Interchromosomal reads between chromosomes 11, 14, derivative 11 and derivative 11 were set to NA.

#### Chrom3D modeling

Chrom3D^30^ was applied to build 100 nuclear models of normal B cells, CLL and MCL. As input the deconvoluted matrices were used, as well as A and B compartment assignments instead of lamina-associated domains. Chrom3D was performed using default settings, with the exception that for each model we shuffled the order in which the chromosomes were fed into the model. Next, per model the nuclear positioning of each chromosome was calculated as the Euclidean distance of its center of mass with respect to the center of the nucleus. In addition, for chromosomes 11 and 14, for each model we calculated the pairwise Euclidean distance per 1 Mb bin.

#### ChimeraX visualization

For visualization of the *in silico*-generated nuclear models, we used ChimeraX (version 1.6). CMM output files of Chrom3D were imported into ChimeraX for whole nuclei visualization. For specific chromosome visualization, the R,G,B values of the CMM files were modified and the files were filtered to display selected beads with desired colors. The modified files were then re-imported into ChimeraX.

### 9. CRISPR-Cas9 genome editing to generate translocations

#### sgRNA design

Single guide RNA molecules were designed to target the region where translocations usually cluster in MCL patients and synthesized by Synthego (Redwood City, CA, USA). The targeted regions of interest in hg38 are chr11:69,531,666-69,531,685 (sequence: GAACCCAGGGTCCATTCCAC) and chr14:105,863,775-105,863,794 (sequence: TCCCTAAGTGGACTCAGAGA).

#### CRISPR-Cas9-based genome editing

Genome editing of the healthy cell line GM12878 to generate translocation t(11;14) *de novo* was conducted using the Neon^TM^ electroporation system. Briefly, RNP complexes were formed by combining 3 μM of each sgRNA and 2 μM SpCas9 NLS recombinant nuclease in R buffer (Neon^TM^ NxT Eletroctroporation System 10-μL Kit, cat. no. N1025, ThermoFisher) for 10-20 min at RT. After RNP complex formation, 0.3 million cells were electroporated using the Neon^TM^ device (Neon^TM^ NxT Electroporation System, NEON1S, ThermoFisher) with 1,600 V, 20 ms and 1 pulse. Cells were immediately transferred to pre-warmed 12-well plates with 1 ml of RPMI-1640 media supplemented with 20% FBS without antibiotics and kept at culture conditions until further analyses were performed.

#### Digital PCR analysis

The presence and quantification of the t(11;14) translocation were assessed using the Digital LightCycler Analyzer System from Roche (Roche, cat. no. 09274804001). Briefly, primers targeting both sides of the breakpoint region were designed, (forward: CGATCTTGCAGTCCTACA; and reverse: TGATGGAGTAACTGAGCC), as well as a probe that hybridizes upstream of the translocation breakpoint in chromosome 14 (AGACACATTCCTCAGCCATCACT / 56-FAM / AGACACATT / ZEN / CCTCAGCCATCACT / 3IABkFQ). Hence, only when the translocation is present, a PCR product is generated, and the fluorophore tag will emit light. Based on a control region targeting chromosome 14 upstream of the breakpoint (forward primer: GAAGCGGAGAGAGGTCAC; reverse primer: TGGCCTTTGCAGCTAATA; and probe sequence: CCAAGTCCGGCCACAGATGTC / 5SUN / CCAAGTCCG / ZEN / GCCACAGATGTC / 3IABkFQ), the fraction of translocated over total alleles was calculated and converted to the fraction of cells carrying translocations (considering that the vast majority of the cells are heterozygous for the translocation). Primers and probes for the digital PCR were purchased from IDT and diluted to a stock concentration of 100 μm upon reception. The dPCR kit and plates used were purchased from Roche (Digital LightCycler 5x DNa Master, cat. no. 09393544001; Digital LC Uni Plate, cat. no. 09033696001).

### 10. Bulk RNA-seq data generation and analysis

#### RNA-seq data of patient data

RNA-seq data from 5 patients and 12 control samples were mined from a previous study^24^. We used the raw and FPKM estimates for downstream analysis.

#### RNA-seq data of genomically engineered cells carrying translocations

Single-stranded RNA-seq data was generated as previously described^54^. Briefly, RNA was extracted and purified using the RNeasy kit for RNA purification (Qiagen, cat. no. 74104), from three biological replicates of GM12878 cell populations containing around 10% of t(11;14) translocated cells, and six GM12878 control samples. Controls 1-3 were electroporated with RNPs containing single guide RNAs (sgRNAs) targeting chromosome 11, while controls 4-6 contained sgRNAs targeting chromosome 14 and Cas9. RNA was purified using the RNeasy kit for RNA purification (Qiagen, cat. no. 74104) and retro-transcribed into cDNA. Then, a qPCR of CCND1 was conducted (primer sequences: CCTGTCCTACTACCGCCTCA and TGGGGTCCATGTTCTGCT) using the Lightcycler 480 (Roche) with SYBR green mix (Roche, cat. no. 04707516001). Next, libraries were prepared for next-generation sequencing by the genomics facility (CRG) and adapter-ligated libraries were amplified and sequenced using 100 bp paired-end reads in a Next-Seq(Illumina). The quality of the raw data was checked using FastQC (v. 0.11.5), and reads were trimmed for low quality and adapter using skewer (v. 0.2.2). The percentage of reads mapping to ribosomal RNA was assessed using riboPicker (v. 0.4.3), and trimmer reads from the 12 samples were mapped to the human Gencode release 46 transcriptome using Salmon (v. 1.5.1). Transcript level estimates were summarized using the R package *tximport* (v. 1.32.0), and raw as well as transcripts-per-million (TPM) counts were used for downstream analysis.

#### Differential gene expression analysis

Differentially expressed genes were defined using the DESeq2 R package (v. 1.44.0)^55^. Briefly, only protein-coding genes expressed in at least one sample (TPM or FPKM values > 1) were included for downstream analysis. Next, the *DESeq* function was used to normalize, estimate dispersion, model fitting and hypothesis testing. For primary samples, the 5 MCL samples were grouped and compared to the 12 healthy mature B-cell samples, merging samples from naive, germinal center, memory and plasma B cells. Only those protein-coding genes with a log2 fold change > 1 in MCL samples compared to all healthy samples and an FDR value lower than 0.1, while being expressed in at least 3 out of the 5 MCL samples, were considered as overexpressed. Downregulated genes were assigned having a log2FC < −1 and an FDR of 0.1. For the genetically engineered cells, translocated cells (Tx1-Tx3) were compared to C1-C6 controls treated with only one sgRNA to exclude the effect of double-strand breaks on chromosomes 11 and 14, and other possible off-target effects. Of note, considering that only 10-12% of the cells are translocated, a 2-fold overexpression in translocated cells would result in a global fold change of 1.1. Therefore, differentially expressed genes were defined as upregulated when having a fold change > 1.1 and an FDR < 0.1 in Tx1-Tx3 compared to C1-C6. PCAs were generated using the *prcomp* function from the state R package (v. 3.5.1), using vst-transformed data.

#### Gene ontology analysis

Gene ontology analyses were performed using the R package EnrichGO from clusterProfiler (v. 4.12.3)^56^, and corroborated using the online enrichR tool^57,58^. All protein-coding genes in the human transcriptome were taken as the background list when compared to global upregulated genes. On specific chromosomal regions (such as chr11 up, chr11 down and chr14 up), only protein-coding genes on the same regions were taken as background.

#### Permutation tests

From all protein-coding genes within the investigated chromosomal segment (chr14 up, chr11 up and chr11 down) a random set of genes was picked, equal in number to the set of upregulated genes. For each 3 Mb bin, the number of random genes was compared to the number of true upregulated genes in that window. This was repeated 10,000 times and p-values were calculated as the frequency in which the number of random genes was higher than the number of upregulated genes plus one, overall divided by 10,000.

### 11. Single-cell RNA-seq data generation and analysis

#### Cell isolation and library preparation

For scRNA-seq analysis, we generated single-cell suspensions of genetically engineered GM12878 cells (samples Tx1-Tx3). Cell partitioning, lysis, barcoding and library generation processes were performed by the IRB genomics facility using the GEXSCOPE Single Cell RNA Library kit (HD) from Singleron (cat. no. 4180031) and following manufacturer’s protocol. Libraries were quantified using Qubit HS and sequenced on an Illumina NovaSeq 6000 S2 platform with an average depth of approximately 200 M reads per cell.

#### Data preprocessing

Raw sequencing reads were processed using the CeleScope (v. 2.0.7 pipeline. Briefly, reads were mapped to the GRCh38 reference genome using the STAR aligner within CeleScope, and gene-level expression matrices were generated. Subsequent analyses were performed using R (v. 4.2.1) and the Seurat package (v. 5.0.1). Cells with high mitochondrial RNA content (>20%), low gene count (<500 or >6,000 genes detected), or low read count (<1,000 or >30,000 reads) were filtered out to remove low-quality or dying cells. Doublets were also identified and removed using scDlbFinder (v. 1.12.0).

#### Data normalization and clustering

Single-cell transcriptomes of the three replicates were combined and log-normalized, and the top 3000 variably expressed genes were identified. Immunoglobulin, HLA, mitochondrial and ribosomal genes were filtered out, and cell cycle phase scores were calculated using the *CellCycleScoring* function from Seurat. Cell cycle phase scores, read counts and percentage of mitochondrial reads were regressed out during data pre-processing to correct for sources of unwanted variation. PCA analysis was performed using the same package. Significant components were selected, and cells were clustered using the Louvain algorithm. Cells were classified based on CCND1 expression (at least one transcript per cell) and visualized using UMAP (performed on the first 30 PCs). Differential genes between CCND1-positive and -negative cells were assigned by the FindMarkers function in Seurat using an FDR threshold of 0.1.

#### Active promoter identification

For each gene, chromatin states in GM12878 were assessed within 1,500 bp up- and downstream of the TSS using previously published data^48^. When active promoter or promoter-associated enhancer marks were present, the promoter was considered active.

## Notes

### Competing Interest Statement

The authors have declared no competing interest.

