## Supplementary figures and images for "Translocations can drive expression changes of multiple genes in regulons covering entire chromosome arms"

### Supplementary Figure 1

##### Supplementary figure 1

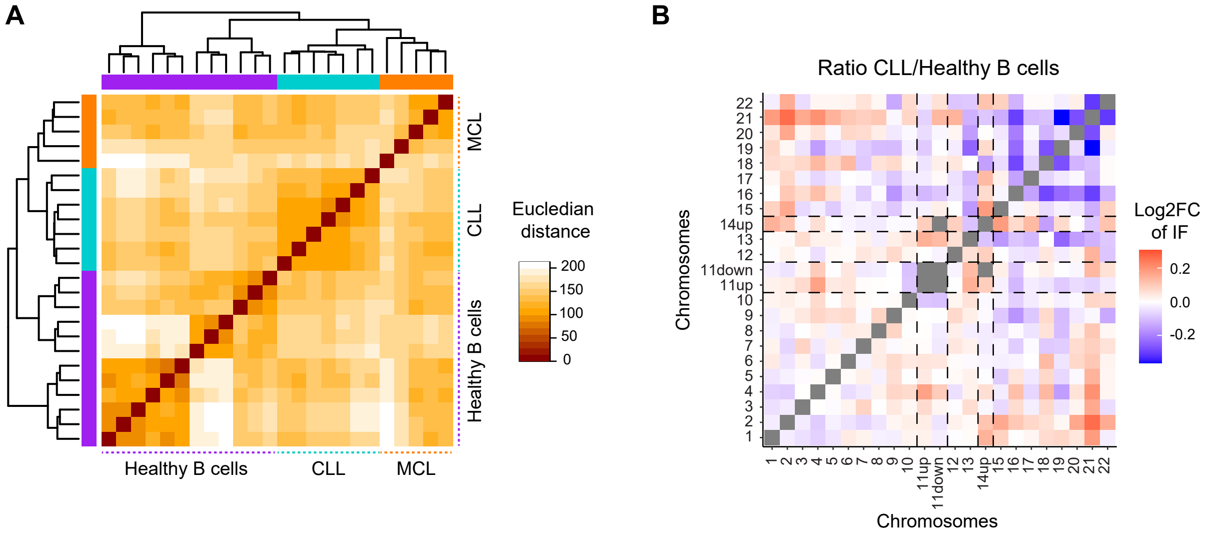

### Supplementary Figure 2

#### Supplementary figure 2

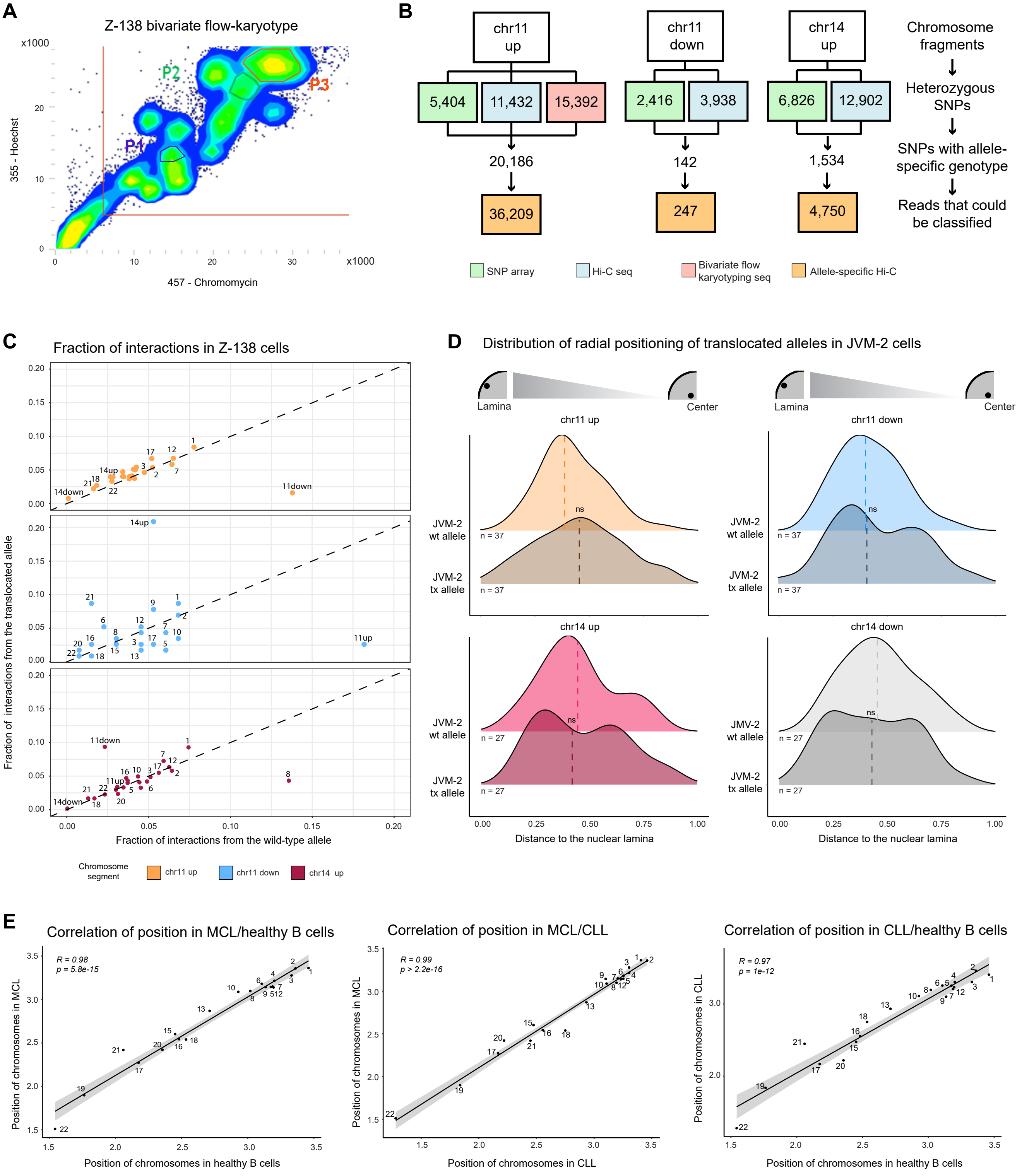

### Supplementary Figure 3

#### Supplementary figure 3

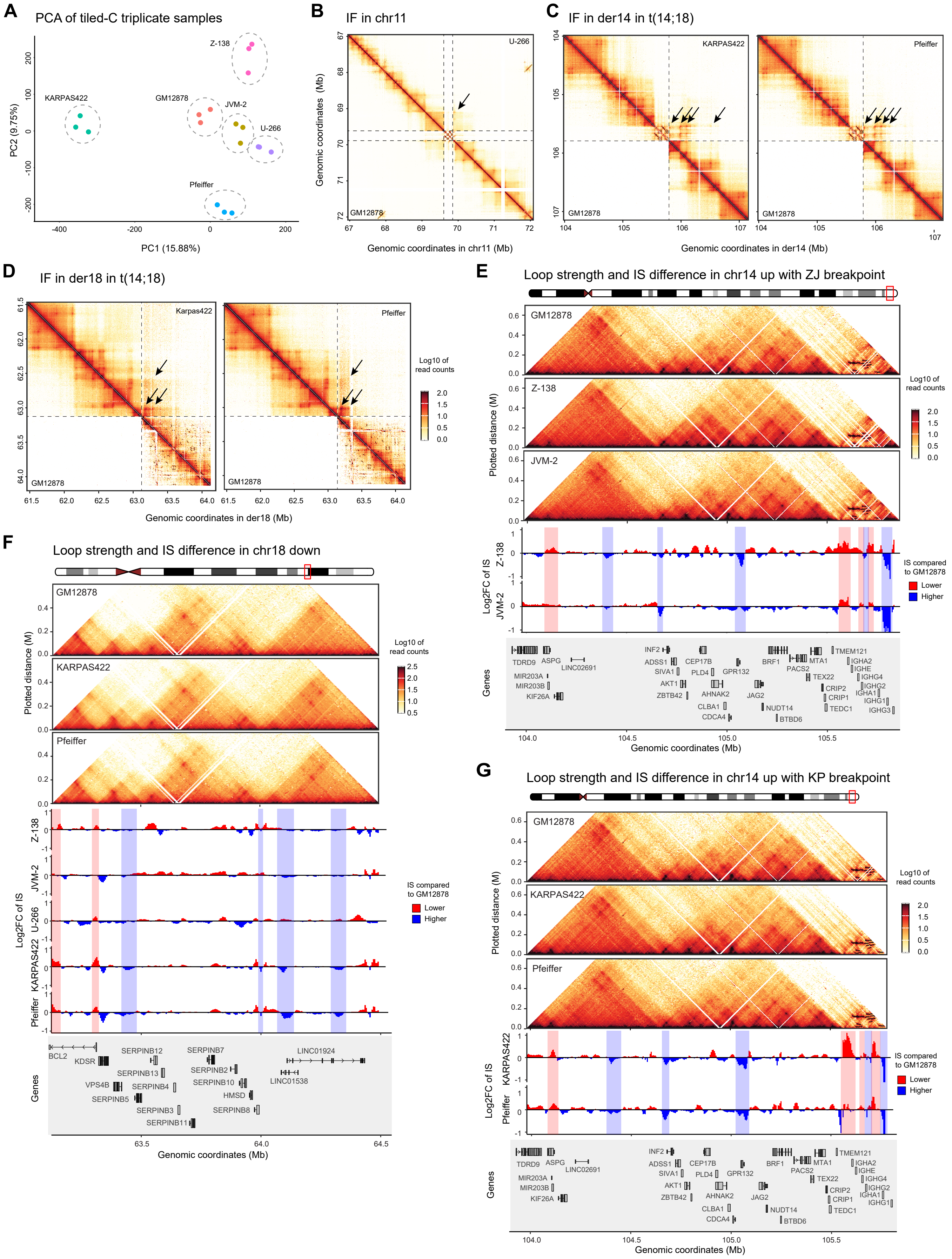

### Supplementary Figure 4

#### Supplementary figure 4

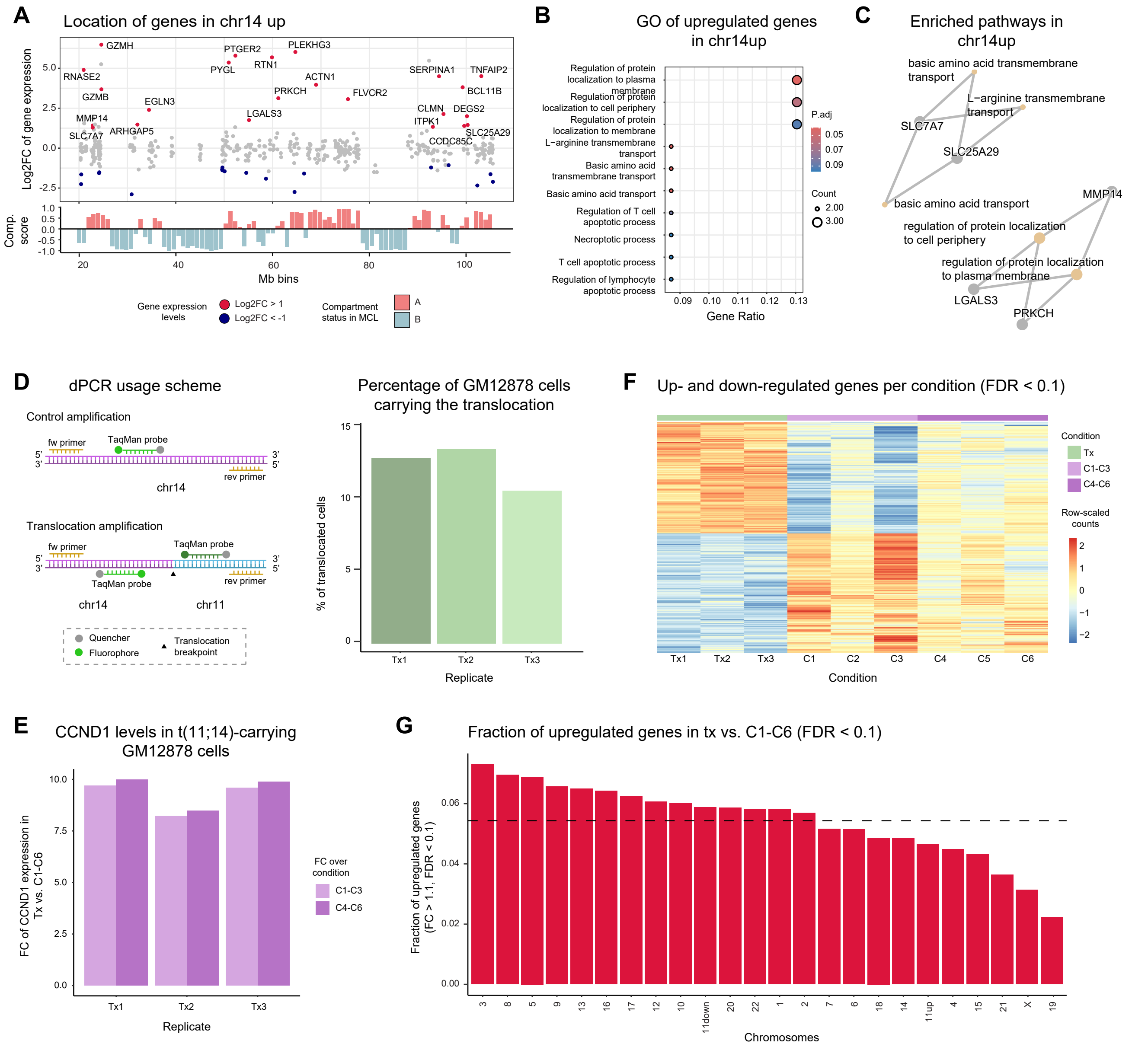
